# Single-Cell Transcriptional Profiling of the Adult Corticospinal Tract Reveals Forelimb and Hindlimb Molecular Specialization

**DOI:** 10.1101/2021.06.02.446653

**Authors:** Noa Golan, Sierra Kauer, Daniel B Ehrlich, Neal Ravindra, David van Dijk, William BJ Cafferty

## Abstract

The corticospinal tract (CST) is refractory to repair after CNS trauma, resulting in chronic debilitating functional motor deficits after spinal cord injury. While novel pro-axon growth activators have stimulated plasticity and regeneration of corticospinal neurons (CSNs) after injury, robust functional recovery remains elusive. These repair strategies are sub-optimal in part due to underexplored molecular heterogeneity within the developing and adult CST. In this study, we combine retrograde CST tracing with single-cell RNA sequencing to build a comprehensive atlas of CSN subtypes. By comparing CSNs to non-spinally projecting neurons in layer Vb, we identify pan-CSN markers including *Wnt7b*. By leveraging retrograde tracing, we are able to compare forelimb and hindlimb projecting CSNs, identifying subtype-specific markers, including *Cacng7* and *Slc16a2* respectively. These markers are expressed in embryonic and neonatal CSNs and can be used to study early postnatal patterning of the CST. Our results provide molecular insight into the differences between anatomically distinct CSN subtypes and provide a resource for future screening and exploitation of these subtypes to repair the damaged CST after injury and disease.

## Introduction

The corticospinal tract (CST) is the major descending motor pathway responsible for fine coordinated movement in mammals.^1^ Corticospinal neurons (CSNs) in layer Vb of sensorimotor cortex project their axons through the internal capsule which then decussate in the brainstem and innervate every spinal segment along the neuroaxis, where they synapse on spinal interneurons whose axons activate motor neurons to initiate and refine movement. The CST is wired postnatally, and after exuberant developmental terminal arborization in spinal grey matter, is pruned via activity-dependent mechanisms during a protracted critical period that ultimately sculpts a mature motor pathway.^2, 3^ The complex wiring of the mature CST, together with its central role in voluntary and fine motor control means that damage leads to significant and lasting functional impairments. Efforts to repair the damaged CST have broadly focused on either nullifying the effects of the axon growth inhibitory environment of the mature CNS, or recapitulating cell autonomous developmental mechanisms to re-build the damaged tract.^4–6^ While strides have been made in stimulating axotomized CSNs to regenerate or intact CSNs to undergo plasticity after injury, recovery of motor function remains woefully incomplete.^7–9^ The inefficacy of these interventions is partially due to an incomplete understanding of the molecular heterogeneity among CSNs. Despite clear anatomical subdivisions of the CST, i.e., CSNs with terminals in the thalamus, brainstem, and cervical and lumbar spinal cord, *in vitro*, and *in vivo* screening approaches designed to identify novel axon growth activators overlook molecular heterogeneity within these neurons and thus dilute the most potent candidates among differentially sensitive subtypes of cells.^8, 10, 11^

Recent data from our laboratory supports differential sensitivity of CSN subdivisions, as we showed that novel pro-plasticity factors identified in intact CSNs undergoing functional plasticity after unilateral pyramidotomy stimulate growth of lesioned and intact forelimb CST axons, while having no effects on lumbar projecting CSNs.^8, 9^ Pro-axon growth candidates *Lppr1* and *Inpp5k* were identified via retrograde labeling of sprouting CSNs in the cervical spinal cord, suggesting that retro-labeling of plastic CSNs from the lumbar would identify a separate set of factors.

Single-cell RNA sequencing (scRNAseq) approaches affords the sensitivity to identify transcriptional heterogeneity among CSNs. Indeed, previous studies have revealed the rich phenotypic diversity among cardinal CNS cell types including neurons, astrocytes, monocytes, macrophages, and microglial cells.^12–19^ Notably, studies in the retina have confirmed molecular heterogeneity within functional subclasses of retinal ganglion cells, enabling the exploration of sub-type specific injury-induced gene expression changes and identifying differences in resilience and axon growth potential after optic nerve injury.^18, 20, 21^ Central to the utility of exploiting transcriptional atlases is understanding the functional role of potential subclasses of neurons. For instance, scRNAseq of cortico-cortico projection neurons showed that while these cells form a single genetic cluster, their anatomically traced subdivisions have diverse gene expression.^22^

To identify transcriptional heterogeneity within the CST, we used an intersectional approach combining retrograde tracing from the cervical and lumbar spinal cord with scRNAseq of adult CSNs. We identified the robust CSN-specific marker *Wnt7b*, and CSN-subtype specific markers including *Cacng7* for forelimb (FL) CSNs, *Slc16a2* for hindlimb (HL) CSNs, and *Rspo2* for dual-projecting CSNs. We believe that these markers can be leveraged for *in vitro* and *in vivo* screening, targeting, and exploitation strategies to enhance our understanding of the development, patterning, housekeeping and response to injury within the CST. CST targeted repair interventions will continue to be functionally sub-optimal until we delineate the molecular heterogeneity within this population.

## Results

### A novel pipeline for scRNAseq of retrogradely labeled CSNs

The CST is the central motor system for controlling voluntary, flexible, and skilled movements. To achieve this, CSNs extend axons from layer Vb of primary motor cortex (M1) to innervate every spinal segment, synapsing principally on spinal interneurons to control FL and HL motor output. Spinal retrograde tracing (**Fig. 1a)** from C6/C7 (**Fig. 1b**) and L4/5 (**Fig. 1c**) show that CSNs that innervate the FL are anatomically distinct from those that innervate the HL (**Fig. 1d, d’**), mirroring functional dexterity differences and suggesting that anatomically distinct CSNs are also molecularly heterogenous. To explore the molecular heterogeneity among CSN populations, we completed scRNAseq on retrogradely labeled FL and HL CSNs and compared gene expression between these anatomically discrete populations to other layer V neurons in M1 in adult mice. As relatively large pyramidal neurons, CSNs have a lower nucleus to cytoplasm ratio, hence a large portion of transcripts are extra-nuclear. Therefore, it was critical to maintain plasma membrane integrity to achieve maximal quality of whole-cell sequencing.^23^ To that end, we refined established adult neuronal dissociation protocols (methods) to develop a novel pipeline for labeling, extracting, dissociating, and sequencing neurons with long-distance axonal projections.^14–16^ To independently label FL and HL CSNs, we microinfused retrogradeAAV-CAG-GFP into the grey matter of either C6/7 or L4/5 spinal cord in adult mice (**Fig. 1e**). After ten days, we macrodissected layer V of M1, enzymatically and physically dissociated the tissue to create a single-cell suspension. Dissociated CSNs were then incubated in a cytoplasmic dye to aid whole cell detection for subsequent fluorescence activated cell sorting (FACS, **supplementary Fig. 1**). Intact CSNs were then collected for scRNAseq using the 10x chromium platform (**Fig. 1e**). Inspection of harvested neurons following dissociation during imaging flow cytometry (Amnis Imagestream, **Fig. 1f)** and after FACS (**Fig. 1g)** clearly shows healthy DAPI positive nuclei surrounded GFP+ cytoplasm. Consistent with robust cellular integrity, sequenced neurons had an average library size of 20,250 unique molecular identifiers (UMIs, median = 19,251 UMIs), and 5,215 genes per cell (median = 5,616).

**Figure 1.**
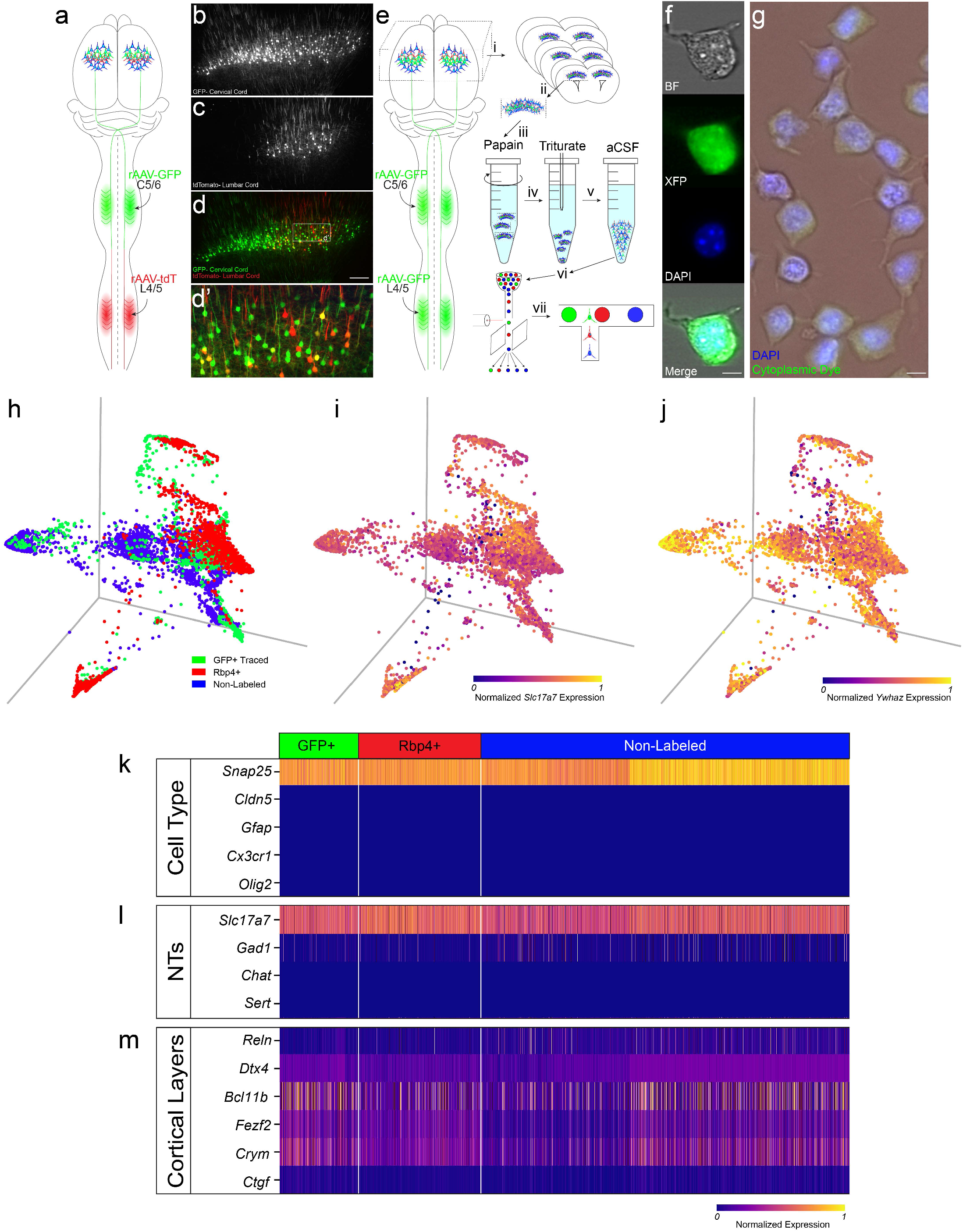
Schematic of spinal retrograde tracing via injecting rAAV-CAG-GFP at spinal level C6/7 and rAAV-CAG-tdTomato at spinal level L4/5 (**a**). Photomicrographs (**b-d**) from primary motor cortex showing FL-CSNs (**b**), HL-CSNs (**c**), and an overlay with both single and dual-projecting CSNs (**d, d’**) 14 days post spinal infusion. Schematic overview of the retroseq procedure (**e**). Mice received retrograde injections of rAAV-CAG-GFP into either C6/7 or L4/5 (i). After a 2-week incubation, M1 was macrodissected (ii), enzymatically dissociated in papain (iii), mechanically dissociated (iv), and incubated in aCSF (v). Traced, *Rbp4*-positive, and nonfluorescent cells were then sorted using FACS (vi) and sequenced using 10x (vii). Cells were inspected for membrane integrity via visualization on an Amnis Imagestream (**f**) and plating after FACS (**g**). PHATE maps colored by cell identity (**h**), *Slc17a7* expression (**i**), and *Ywhaz* expression (**j**), further confirming cellular integrity. Heatmaps representing cell-type expression (**k**), neurotransmitter expression (**l**), and cortical layer marker expression (**m**), confirming that all sequenced cells are layer V, glutamatergic neurons.

Previous studies have detailed molecular differences between and among excitatory and inhibitory neurons spanning the entire cortical depth.^12, 14–16, 24–27^ Here we focused specifically on molecular differences among layer V neurons and comprehensively characterize transcriptional specificity between CSNs. To achieve this, we compared gene expression between retrogradely traced CSNs, layer Vb non-fluorescent neurons (controls), and neurons expressing the previously identified layer-V marker *Rbp4*, sequencing a total of 5793 adult neurons (**Fig. 1e**).*^15^* To explore gene expression differences among these populations, we used potential of heat diffusion for affinity-based transition embedding (PHATE) for dimensionality reduction and visualization, as unlike PCA, t-SNE and uMAP, PHATE maintains both local and global structure, and was designed for scRNAseq (**supplementary Fig.2)**.^28, 29^ PHATE maps labeled with cell origin (**Fig. 1h**) and *Slc17a7* expression (**Fig. 1i)** demonstrate that while all sequenced cells are excitatory pyramidal neurons, they exist on a continuous spectrum of cellular phenotypes, rather than discrete cell types. Critically, gene expression visualization with PHATE further confirms the cytoplasmic integrity of sequenced neurons as they are all enriched in the whole-neuron-specific marker *Ywhaz* **(Fig. 1j**).^26^

To broadly confirm identity of sequenced cells, we assessed their expression of previously identified cell-type specific markers. All sequenced cells expressed the neuron-specific marker *Snap25* and showed a dearth of expression for other cell-type markers including the endothelial marker *Cldn5*, the astrocyte marker *Gfap*, the microglial marker *Cx3cr1,* and the oligodendrocyte specific marker *Olig2* (**Fig. 1k**). Neurotransmitter gene expression confirmed that all sequenced neurons expressed the glutamatergic marker *Slc17a7* and not the GABAergic marker *Gad1*, the cholinergic marker *Chat* or the serotonergic marker *Sert* (**Fig. 1l**). All sequenced neurons showed a moderate enrichment for established layer V markers *Bcl11b, Fezf2,* and *Crym*, relative to other cortical layer makers *Reln, Dtx4*, and *Ctgf* (**Fig. 1m**), confirming layer V identity.^12, 30^ While CSNs expressed previously identified layer V markers, none of those markers were specific to any layer V population, suggesting that more targeted analyses are necessary to identify unique molecular characteristics of CSNs.

### Identifying CSN-specific markers in Layer V of M1

To identify CSN specific makers within M1, we compared traced neurons to layer Vb non-fluorescent control neurons (**Fig. 2a**). Differential expression (DE) analysis revealed 565 CSN-specific differentially upregulated genes and 695 downregulated genes (**Fig. 2b**). We used recursive feature elimination to determine the ten upregulated and downregulated genes most strongly differentiated between CSNs and control cells. The top genes that define CSNs include *Wnt7b, Slc5a5,* and *S100b,* while the top genes that are enriched in adjacent layer Vb control cells include *Smoc2, Calb1* and *Cux2* (**Fig. 2c**). To validate these data, we trained a linear classifier to predict cell identity based on sole expression of either the top ten upregulated CSN genes, or the expression of eleven genes that have previously been used to describe layer V pyramidal neurons including *Npsr1, Fezf2, Bcl11b, Crym,* and *Colgalt2*.^12, 15, 16, 30–32^ Analysis of the accuracy and precision of the classifier performance using these groups of genes shows that our new list of CSN-specific genes are significantly better predictors of CSN identity (**Fig. 2d**).

**Figure 2.**
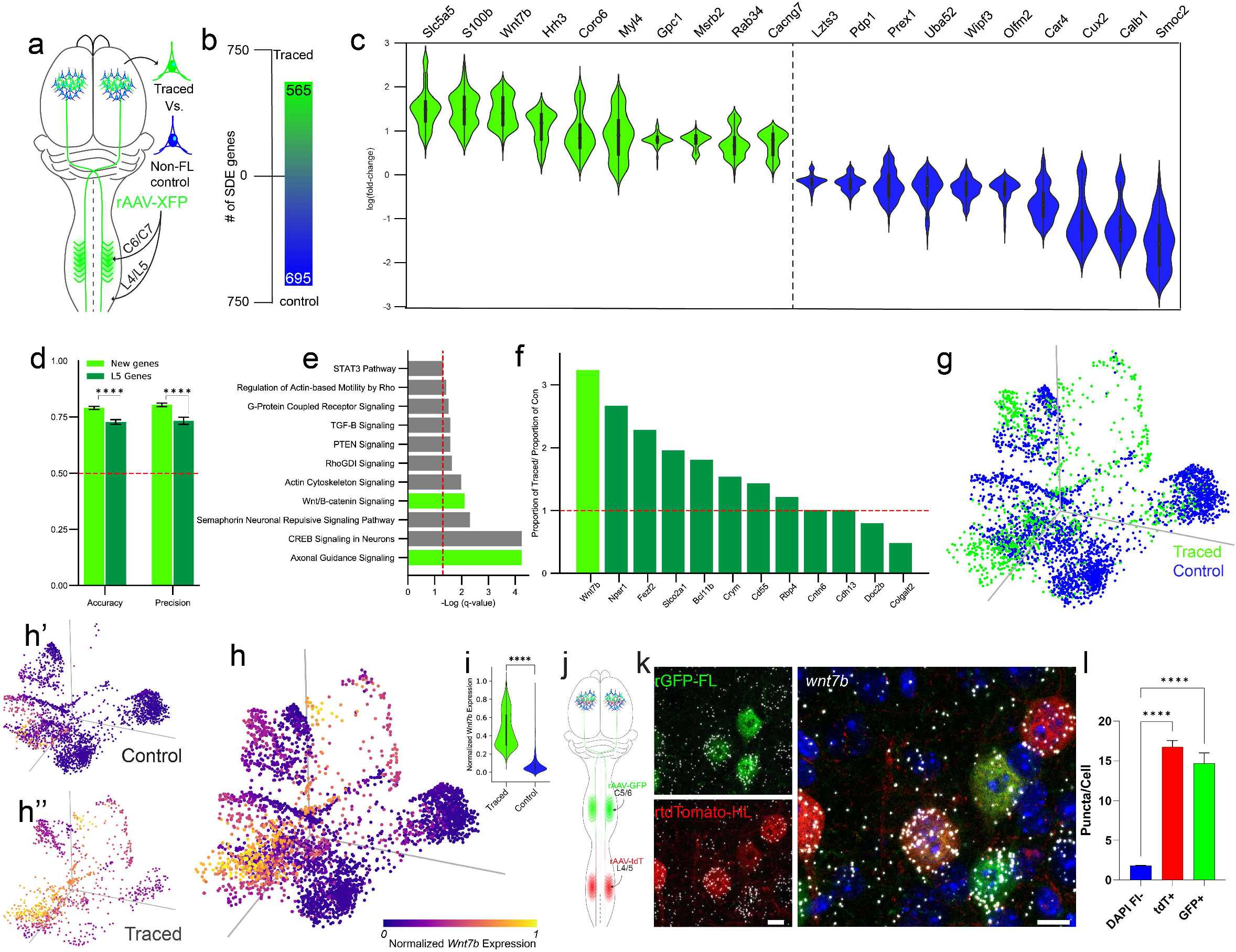
Schematic overview of retrogradely labeled CSNs compared to nonfluorescent layer V control neurons (**a**). Differential expression analysis identified 566 upregulated genes in CSNs and 696 downregulated genes vs. control neurons (**b**). Violin plots of the top 10 differentially expressed genes in traced CSNs and control neurons (**c,** Wilcoxon rank sum test with a Benjamini-Hochberg correction, p-adj < 0.05). Accuracy and precision of a linear support vector machine (l-svm) in prediction of CSN identity is significantly higher based on the top 10 DE genes, light green, compared to previously identified layer V genes, dark green (**d,** Wilcoxon rank sum test with a Benjamini-Hochberg correction, p < 0.001). Ingenuity pathway analysis showing significantly enriched pathways associated with DE genes. Red line denotes significance of q<0.05 (**e**). *Wnt7b* is expressed in a higher proportion of CSNs relative to controls compared to other previously characterized layer V markers. Red line denotes equal expression in CSNs and control neurons (**f**). PHATE maps colored by cell identity (**g**) and *Wnt7b* expression in just control neurons (**h’**), just traced neurons (**h”**) and all neurons (**h**) illustrate *Wnt7b* CSN enrichment. *Wnt7b* is significantly enriched in traced compared to control cells (**i**, Wilcoxon rank sum test with a Benjamini-Hochberg correction, p-adj < 0.001). Schematic showing retrograde labeling of cervical CSNs (GFP) and lumbar CSNs (tdTomato) (**j**). smFISH of retrogradely labeled CSNs shows an enrichment of *Wnt7b* in CSNs compared to nonfluorescent, DAPI cells in layer Vb (**l**, data shown are average number of puncta per cell ± SEM, One-way ANOVA with a Bonferroni correction, p < 0.0001). Scale bar, k = 10um.

To explore the biological import of the transcriptional differences that define CSNs, we completed Ingenuity pathway analysis (IPA).^8^ Amongst the top enriched pathways were those associated with cytoskeletal dynamics, including actin cytoskeleton signaling, regulation of actin-based motility by rho, Rho GDI signaling; axon growth, including STAT3, PTEN and CREB signaling; and pathways associated with axon guidance including semaphorin neuronal repulsive signaling pathway, and Wnt/β-catenin signaling (**fig. 2e**). Axon guidance cues including *Epha4, Efnb3, Ryk* and *Wnt3a* have previously been shown to be critical in wiring the CST. ^33^ Based on these observations, we selected *Wnt7b* as a primary candidate to validate our novel CSN specific marker gene list.

### Validation of *Wnt7b* as a CSN-specific marker

DE analysis revealed that *Wnt7b* was significantly enriched in CSNs. Evaluation of the proportion of traced versus non-traced cells showed that *Wnt7b* was expressed in a 3:1 ratio, outperforming all previously described layer V markers (**Fig. 2f, supplementary Fig. 3**). This analysis reveals that many established layer V marker genes are not CSN-specific and some, including *colgalt2* and *doc2b* are expressed in a higher proportion of non-spinally projecting layer V control cells. PHATE visualization of all layer V cells sequenced allows us to evaluate *Wnt7b* expression in traced and control cells independently (**Figs. 2g, h**). Both traced and control cells occupy relatively similar space in PHATE (**Fig. 2g**). However, heat maps of *Wnt7b* expression per cell of origin illustrates robust difference between CSNs and control cells (**Fig. 2h**). These data confirm that *Wnt7b* is significantly enriched in all traced cells (**Fig. 2i)**. To validate this finding *in vivo,* we infused rAAV-CAG-GFP into the grey matter of C6/C7 spinal cord and rAAV-CAG-tdTomato into the grey matter of L4/L5 spinal cord in adult wild type mice, differentially labeling FL and HL CSNs and completing single molecule fluorescent *in situ* hybridization (smFISH, **Fig. 2j**). Photomicrographs of Layer Vb from M1 prepared 2 weeks after spinal infusion show GFP+ FL CSNs and tdTomato+ HL CSNs (**Fig. 2k**) are significantly enriched in *Wnt7b* relative to DAPI+ control cells (**Fig. 2l**).

### Molecular identification of anatomically distinct CSN subtypes

While CSNs can be defined by the adult expression of *Wnt7b*, we next wanted to explore the potential molecular heterogeneity between anatomically discrete FL and HL projecting CSNs. PHATE visualization of traced neurons colored by anatomical origin illustrates that there is a clear transcriptional separation between FL and HL traced cells (**Fig. 3a, b**). DE analysis revealed that there were 737 FL-specific genes and 492 HL-specific genes (**Fig. 3c**). Identifying a set of anatomically specific FL and HL CST markers is central to understanding the potential independent homeostatic biology and injury-induced changes within these discrete populations. To prioritize candidate markers, recursive feature elimination was used to determine the top ten FL- and HL-specific genes that most robustly distinguished these two populations. The top genes that define the FL population include calcium related genes *Cacng7* and *Pcp4,* as well as the structural gene *Sgcz* (**Fig. 3d, supplementary Fig. 4**). The top genes that define the HL population include thyroid transporter *Slc16a2* and the transcriptional repressors *Bcl6* and *Bcorl1* (**Fig. 3d, supplementary Fig. 5**).

**Figure 3.**
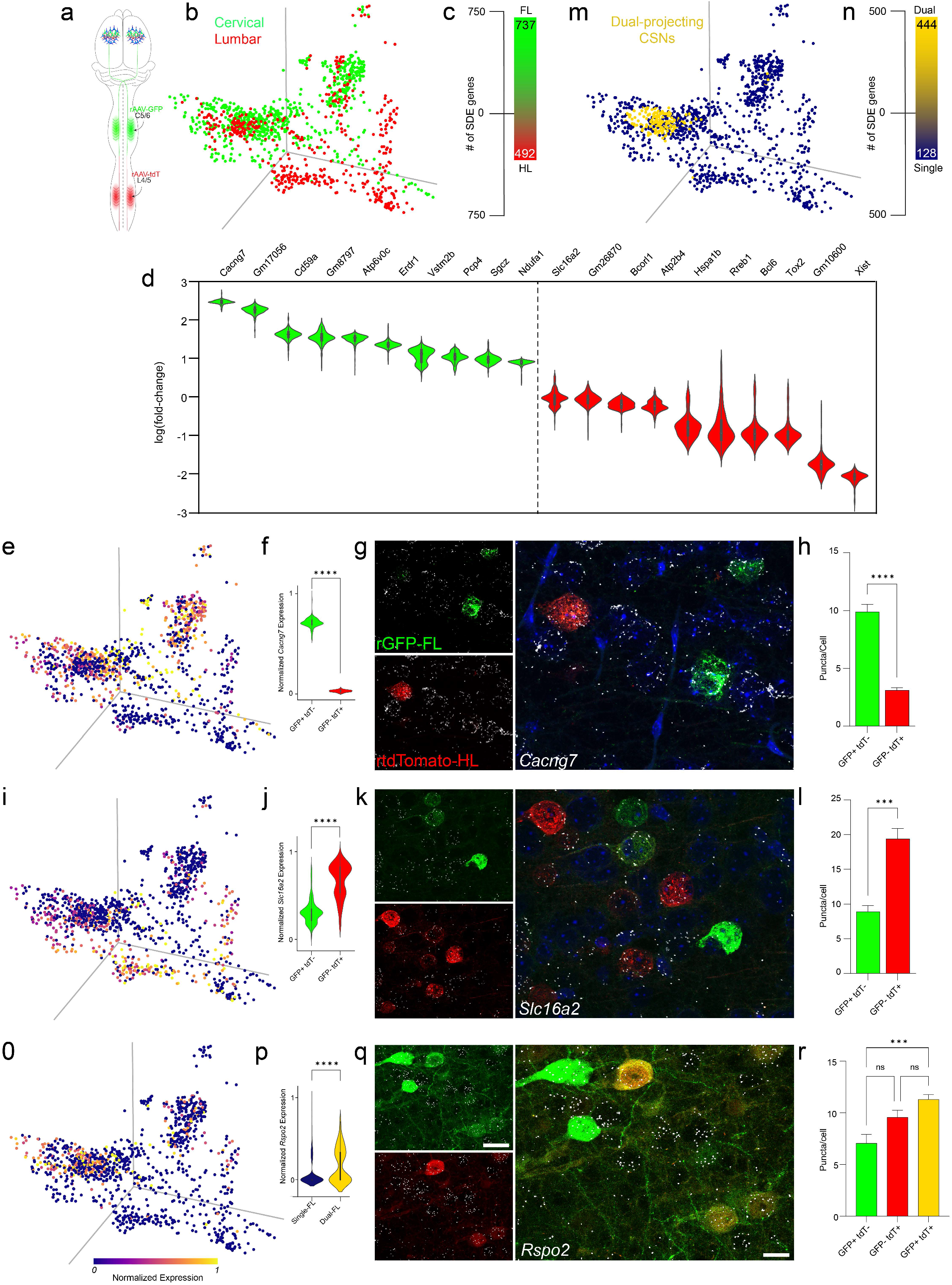
Schematic of dual spinal retrograde labeling paradigm (**a**). PHATE map colored by cell identity of retrogradely labeled GFP+ cervical CSNs and tdTomato+ lumbar CSNs shows a broad separation (**b**). Differential expression analysis identified 738 upregulated FL genes and 492 upregulated HL genes (**c**). Violin plots of the top 10 highest differentially expressed genes in FL CSNs (green) and HL CSNs (**d**, red, Wilcoxon rank sum test with a Benjamini-Hochberg correction, p-adj < 0.05). PHATE map colored by *Cacng7* expression illustrates FL enrichment (**e,** compare to **b**). *Cacng7* is significantly enriched in FL compared to HL cells (**f**, Wilcoxon rank sum test with a Benjamini-Hochberg correction, p-adj < 0.001). smFISH of retrogradely labeled GFP+ FL-CSNs and tdTomato+ HL-CSNs shows an enrichment of *Cacng7* in FL-CSNs (**g**, **h**, data shown are average number of puncta per cell ± SEM, unpaired t-test, p < 0.0001). PHATE map colored by *Slc16a2* expression illustrates HL enrichment (**i**, compare to **b**). *Slc16a2* is significantly enriched in HL-CSNs compared to FL-CSNs (**j**, Wilcoxon rank sum test with a Benjamini-Hochberg correction, p-adj < 0.001). smFISH of retrogradely labeled GFP+ FL-CSNs and tdTomato+ HL-CSNs shows an enrichment of *Slc16a2* in HL-CSNs (**k**, **l**, data shown are average number of puncta per cell ± SEM, unpaired t-test, p < 0.001). PHATE map showing dual FL and HL projecting CSNs in yellow (**m**). Differential expression analysis identified 738 upregulated FL genes and 492 upregulated HL genes (**n**). PHATE map colored by *Rspo2* expression illustrates dual enrichment (**o**). *Rspo2* is significantly enriched in dual-projecting CSNs compared to FL and HL CSNs (**p**, Wilcoxon rank sum test with a Benjamini-Hochberg correction, p-adj < 0.001). smFISH of retrogradely labeled GFP+ FL-CSNs and tdTomato+ HL-CSNs shows an enrichment of *Rspo2* in dual projecting CSNs relative to FL CSNs (**q, r**, data shown are average number of puncta per cell ± SEM, one-way ANOVA with a Bonferroni correction, p < 0.001) and a trend of enrichment relative to HL-CSNs (**r**, data shown are average number of puncta per cell ± SEM one-way ANOVA with a Bonferroni correction, p = 0.06).

PHATE visualization of *Cacng7* illustrates significantly enriched expression in FL traced neurons (**Fig. 3e, f**). To validate these results *in vivo*, mice received injections of rAAV-CAG-GFP into spinal grey matter at C6/7 and rAAV-CAG-tdTomato into spinal grey matter at L4/5. Visualization of *Cacng7* using smFISH confirmed differential expression in GFP-positive FL-projecting neurons compared to tdTomato-positive HL-projecting neurons (**Fig. 3g, h**). A similar analysis was then repeated with the HL-specific marker *Slc16a2* which was significantly enriched in HL-projecting neurons (**Fig. 3i, j**). *In vivo* smFISH confirmed HL-specificity with enrichment of *Slc16a2* in tdTomato-positive HL-projecting neurons relative to GFP-positive FL-projecting neurons (**Fig. 3k, l**).

### Identification of dual FL-HL projecting CSNs

Histological analysis of spinal retrograde tracing from C6/C7 (**Fig. 1b**) and L4/5 (**Fig. 1c**) shows that there is a population of neurons that express both tracers and therefore has terminals in both cervical and lumbar cord (**Fig. 1d**). These observations are confirmed transcriptionally as PHATE mapping of all traced cells shows small overlapping regions between these populations (**Fig. 3a**). As tracing and sequencing of CSNs from cervical and lumbar targets was completed in independent mice, we hypothesized that cells present in overlapping regions in PHATE correspond to dual-projecting CSNs. We used kernel density estimation to identify FL cells that had a high probability of also being HL cells and HL cells that had a high probability of also being FL cells (**supplementary Fig. 6**). This analysis identified a cluster of dual-projecting neurons (**Fig. 3m**). Differential expression analysis between dual-projecting CSNs and the single projecting CSNs identified 444 significantly upregulated genes and 128 downregulated genes (**Fig. 3n**). Of these genes *Rspo2*, an enhancer of Wnt signaling, was one of the top DE genes. To validate, PHATE visualization of *Rspo2* expression confirms dual-projecting specificity (**Fig. 3o, p**). *In vivo* validation with smFISH shows an enrichment of *Rspo2* expression in dual-projecting tdTomato- and GFP-positive neurons relative to GFP-positive FL-projecting neurons and a trend towards enrichment when compared to tdTomato-positive HL-projecting neurons (**Fig. 3q, r**).

### Dorsoventral specificity of CSN terminals

To explore whether CSNs can also be spatially defined by their dorsoventral terminal pattern in spinal grey matter, we completed a ligand-receptor interaction analysis between our HL-CSN scRNAseq data and single-nucleus RNAseq data of lumbar spinal interneurons (SINs).^13^ Using the CellCellInteractions (CCI) dataset, we narrowed down receptor-ligand pairs that were significantly differentially upregulated in traced CSNs and whose cognate pairs were expressed in lumbar SINs.^17^ An expression index was calculated per lumbar interneuron cluster by dividing the number of genes from the narrowed down CCI dataset that were expressed by the total number of genes expressed and a one-tailed permutation test was used to calculate the significance of the expression index for each cluster. This identified nine significant lumbar SIN subtypes that are the likely postsynaptic partners of HL-CSNs: 8 dorsal clusters and one medial cluster (**Fig. 4a**). These data suggest a predominant dorsal innervation pattern of the CST in the lumbar spinal cord. To confirm this, we injected AAV1-tdTomato unilaterally into HL motor cortex in adult wild type mice. Transverse section through L4 prepared 14 days post viral infusion showed that the significantly more CST axons terminated in the dorsal horn (**Fig. 4b, c**). To further classify interactions unique to this circuit, we completed a differential gene expression analysis between the nine identified lumbar SINs clusters and the remaining clusters. Ligand/receptor pairs that were differentially upregulated in both HL-CSNs and the lumbar SINs were identified (**Fig. 4d**). These data show that adult CSNs can be broadly defined by their terminal innervation both the dorsoventral and rostrocaudal spinal axis. Furthermore, this level of terminal specificity provides a transcriptional blueprint for rewiring the CST after injury.

**Figure 4.**
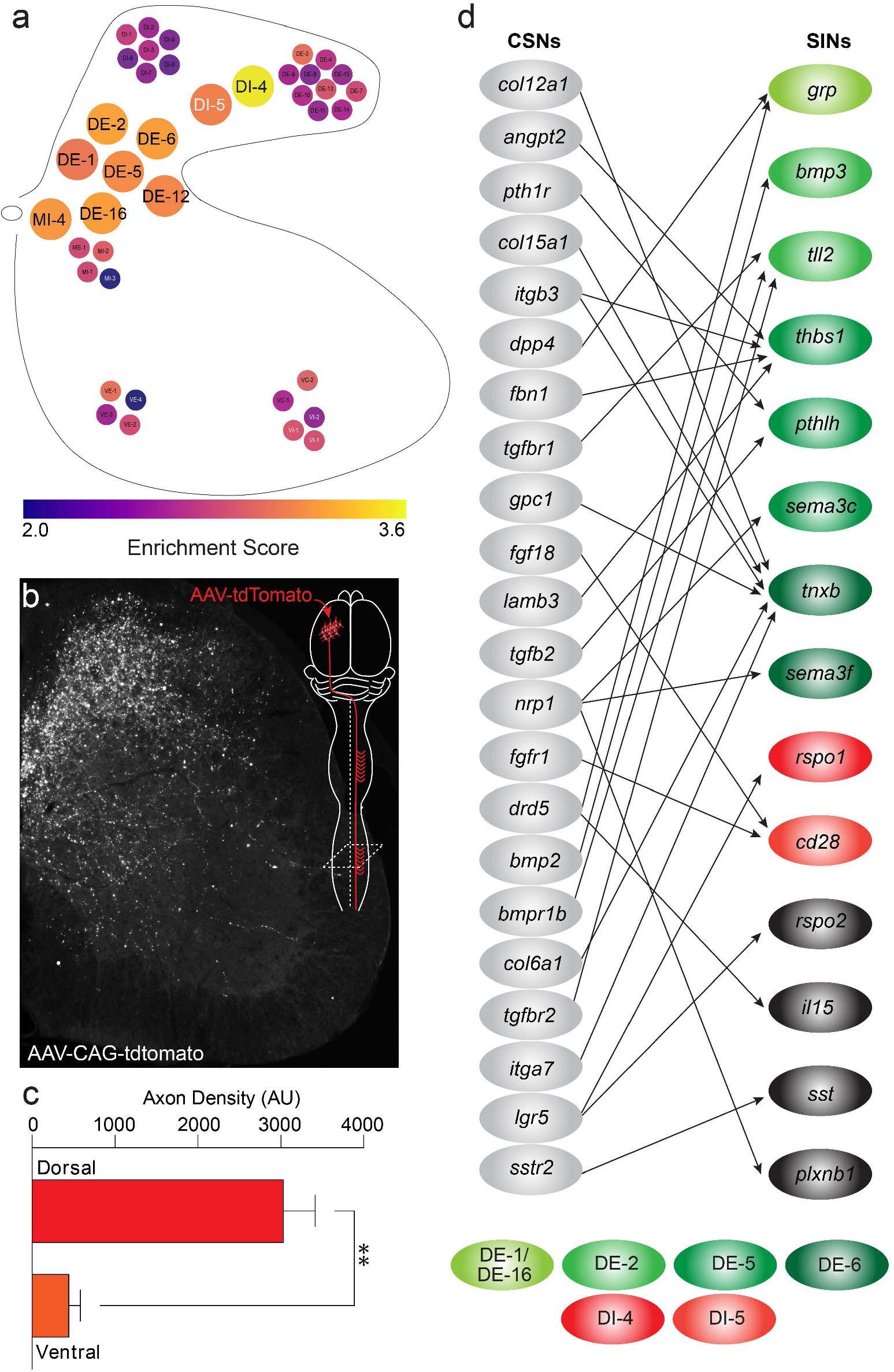
Schematic of the lumbar spinal grey matter showing SIN subtypes identified by Sathyamurthy *et al*., SINs are color coded by enrichment score calculated via ligand/receptor pairs enriched between HL-CSNs and lumbar SINs, 9 enlarged cells receive significant CSN input (**a**, p< 0.05, one-tailed permutation test). Photomicrograph of a lumbar spinal cord 14 days after anterograde tracing with AAV1-tdTomato shows significantly more CST terminals in the dorsal and intermediate zones (**b**), confirmed with densitometric analysis (**c**). Differential gene expression analysis identified a number of ligand receptor pairs that were enriched in these 9 subtypes compared to non-enriched SINs. Genes are colored by enrichment in individual SIN clusters, with the four genes that are enriched in multiple SINs and therefore are uncolored (**d**).

### Leveraging CSN specificity during postnatal development *in vivo*

Central to the utility of exploiting these data to understand the molecular mechanisms that drive wiring of the CST during postnatal development, and re-wiring of the CST after injury, will be our ability to follow these subpopulations during all phases of CST development. To determine whether all CSNs and those innervating the FL and HL can be identified early postnatally, we harvested cortical neurons from *Rbp4 cre:*Ai14 mice at postnatal day (P)5. We selected P5 as at this timepoint, FL and HL-projecting CSNs are in different phases of growth. FL-projecting CSNs have reached the cervical cord and are engaged in local terminal plasticity, while most HL-projecting CSNs are still engaged in long-distance axon growth.^2^ To explore the gene expression profile within layer V at P5, we completed scRNAseq of macro-dissected P5 cortical neurons guided by visualizing layer V via expression of tdTomato in *Rbp4*+ neurons. PHATE maps show that all sequenced layer V neurons (n=2,720) were enriched in *Slc17a2* (**Fig. 5a**) and *Ywhaz* (**Fig. 5b**) demonstrating that were excitatory neurons with intact somata. As expected, only a fraction of cells sequenced were enriched in *Wnt7b* (**Fig. 5c**, 28.1%), consistent with their CSN identity. Exploring gene expression within *Wnt7b*+ neurons (**Fig. 5d**) revealed an enrichment in the CST-specific axon guidance cue *Epha4* (**Fig. 5e**), and the growth associated protein *Gap43* (**Fig. 5f**), confirming these cells as CSNs and demonstrating that they are uniformly in an active growth mode.^34^ *Wnt7b*-positive CSNs separated into two clusters. Remarkably, these clusters are defined by high *Cacng7 (***Fig. 5g)** and *Slc16a2* (**Fig. 5h**) expression, thereby demonstrating that at P5 FL and HL projecting CSN can be transcriptionally defined and thus independently interrogated by temporal gene expression analysis to explore the molecular mechanisms that drive anatomically discrete patterning of the CST.

**Figure 5.**
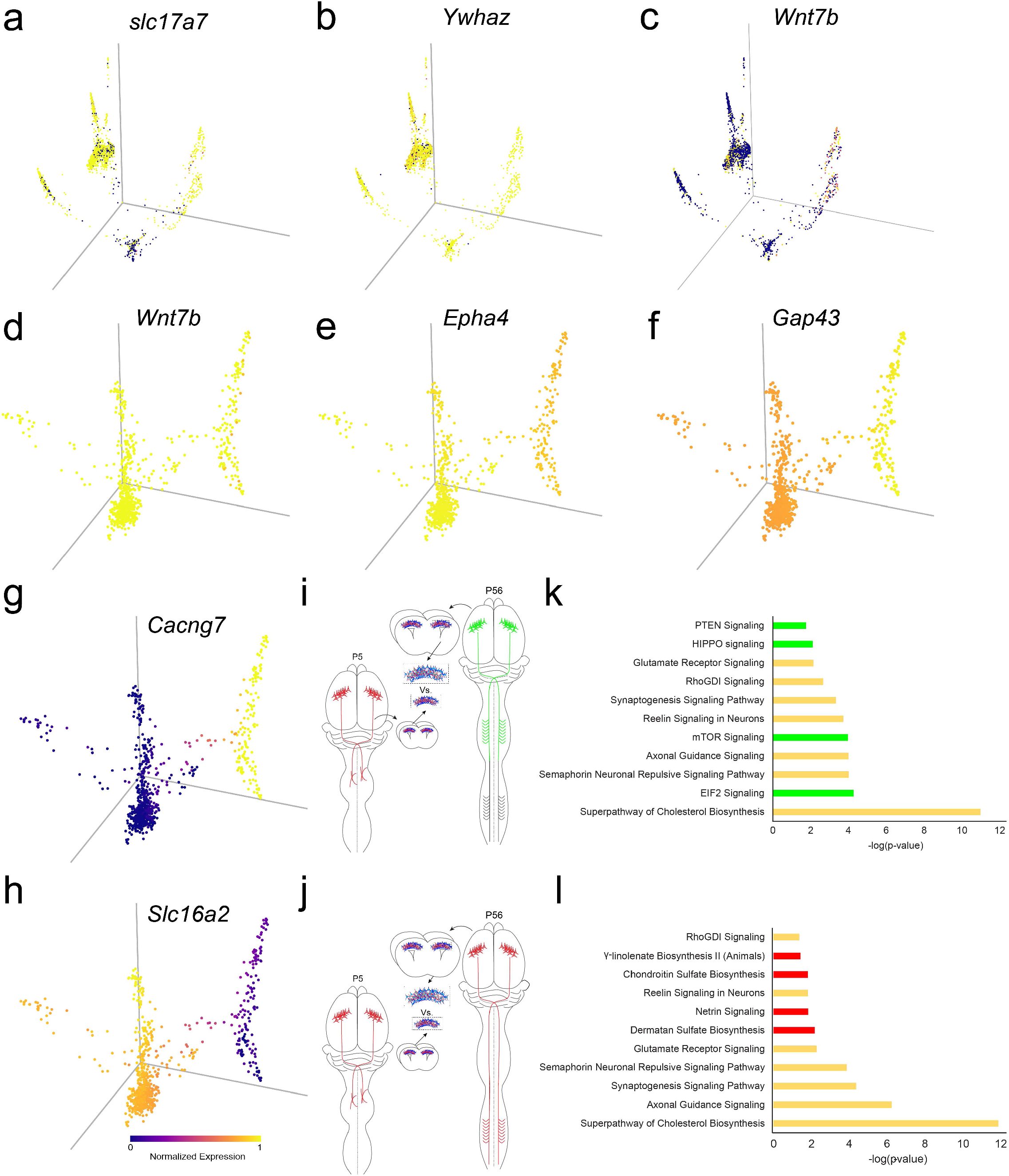
PHATE plots showing that all P5 sequenced neurons express the excitatory marker *Slc17a7* (**a**), the whole cell cytoplasmic marker *Ywhaz* (**b**), and that only a subset of P5 neurons express the CSN marker *Wnt7b* (**c**). PHATE plots confirming P5 CSN identity showing imputed expression of pan-CSN markers *Wnt7b* (**d**), CSN-specific guidance gene *Epha4* (**e**), and the axon growth marker *Gap43* (**f**). FL CSN identity is confirmed via imputed *Cacng7* expression (**g**) and HL CSN identity is confirmed via imputed *Slc16a2* expression (**h**). To identify developmental CSN genes, a differential expression analysis comparing P5 *Cacng7*+ CSNs to adult FL CSNs (**i**) and comparing P5 *Slc16a2*+ CSNs to adult HL CSNs (**j**) was performed. Differentially expressed genes from both of these analyses were enriched in pathways associated with axon guidance and elongation (yellow) as well as FL-specific pathways associated with local plasticity (green, **k**) and HL-specific pathways associated with axon guidance (red, **l**).

To initially surveil the transcriptional machinery that is driving axon growth within these two sub-populations, we used DE analysis to compare P5 FL-CSNs with P56 FL-CSNs (**Fig. 5i)** and P5 HL-CSNs with P56 HL-CSNs (**Fig. 5j**) followed by IPA on the upregulated genes at P5. We found that both FL and HL P5 CSNs show an enrichment for pathways associated with growth, including pathways involved in cholesterol synthesis consistent with axon elongation, as well as pathways involved in axon guidance and synaptogenesis (**Fig. 5k, l**). However, some pathways were unique to one subtype. Pathways specifically enriched at P5 in FL CSNs included those previously associated with axonal sprouting and plasticity including PTEN signaling, Hippo signaling, mTOR signaling, and EIF2 signaling (**Fig. 5k**).^7, 8, 35^ Pathways specifically enriched at P5 in HL projecting CSNs included those associated with axon guidance and neuronal interactions with the extracellular matrix including netrin signaling, y-linolenate biosynthesis, chondroitin sulfate biosynthesis, and dermatan sulfate biosynthesis (**Fig. 5l**).^36, 37^ These data show that FL and HL CSNs have unique molecular profiles at P5 which may reflect the different growth phases between terminal sprouting (FL) versus long distance axon growth (HL). A more acute temporal investigation of gene expression within these two populations will resolve the molecular machinery that is required for these growth phases, which can then be exploited for targeted repair post injury. These profiling studies will only be possible by leveraging *Wnt7b*, *Cacng7* and *Slc16a2* as specific CSN markers.

### Leveraging CSN specificity *in vitro*

Confirming that FL and HL CSNs can be transcriptionally defined at P5 suggests that these markers can also be leveraged to refine functional screens that utilize embryonic cortical neurons *in vitro*.^38^ To explore, we evaluated the expression of *Wnt7b*, *Cacng7*, and *Slc16a2* in embryonic day 17 (E17) dissociated cortical neuron cultures and assessed the neurite outgrowth signatures between these defined populations. At 8 days *in vitro* (DIV), cultures were fixed and stained with TUJ1 and underwent processing with our marker smFISH probes.

We found that *Wnt7b* was enriched in cortical neurons with a neurite length greater than 200um (**Fig. 6a, b**). These data are in line with the expectation that spinally projecting CSNs will support long distance axon/neurite growth. As *Cacng7* and *Slc16a2* are CSN-subtype specific, we limited our analysis of these genes to neurons with processes longer than 200um. Analysis of *Cacng7* revealed no relationship between neurite length and gene expression (**Fig. 6c, d**); however, *Slc16a2* was enriched in neurons with neurite length greater than 300um (**Fig. 6e, f**). These analyses show that CSN-specific markers are expressed at E17 and can be used to identify CSNs *in vitro*, and thus can be leveraged to refine pro-axon growth screening approaches to specifically target all CSNs or those that innervate the FL and HL separately.

**Figure 6.**
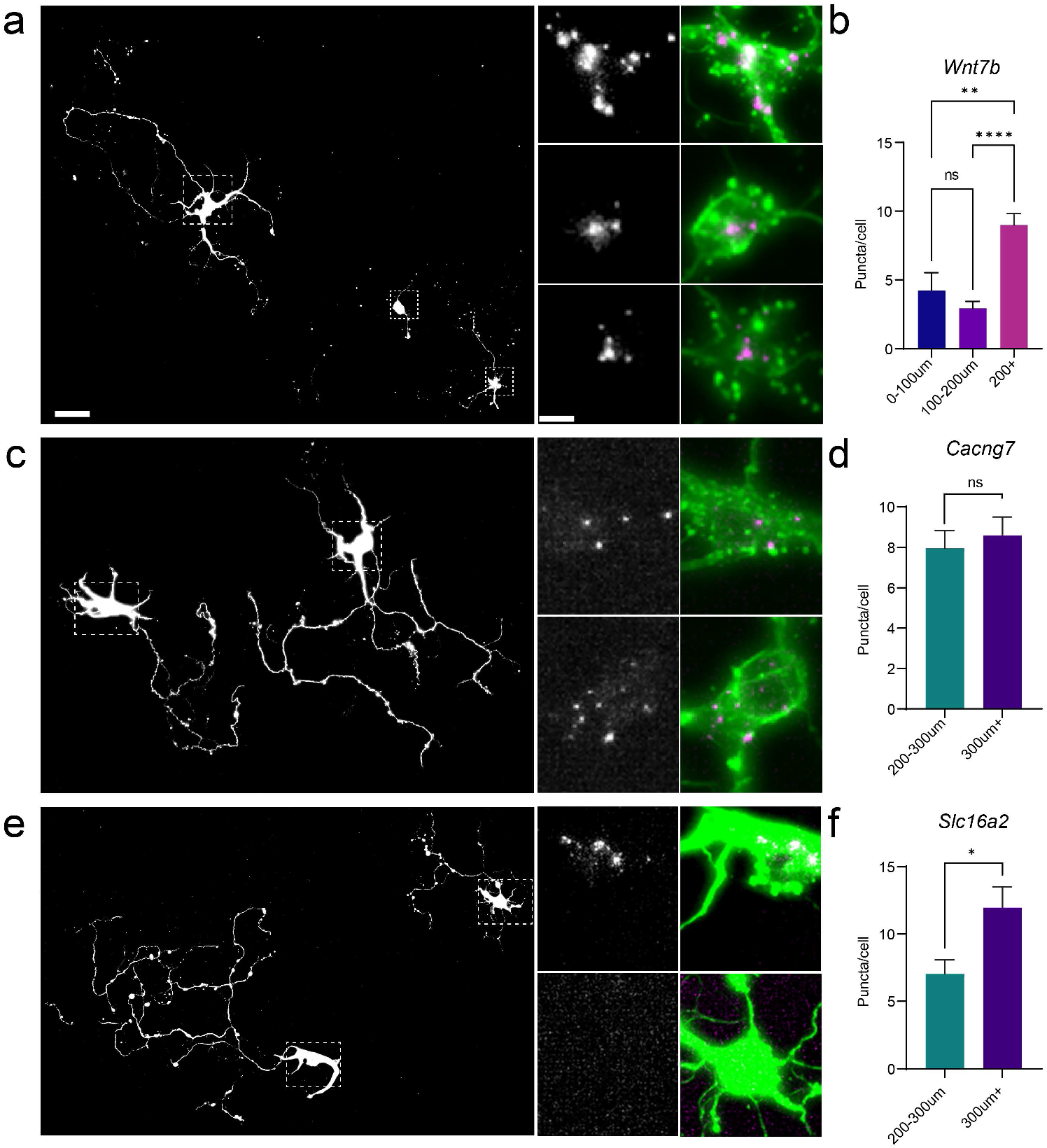
Photomicrographs of E17 cortical neurons 8 DIV stained with β-tubulin. smFISH for *Wnt7b* **(a**) shows enrichment of mRNA in neurons with neurites longer than 200 um (**b**, data shown are average length of neurites per cell ± SEM one-way ANOVA with a Bonferroni correction, ** p < 0.01, **** p<0.0001). smFISH for *Cacng7* (**c**) shows consistent expression in neurons with processes longer than 200um (**d**, data shown are average length of neurites per cell ± SEM independent t-test, p = 0.6). smFISH for *Slc16a2* (**e**) shows enrichment in neurons with processes longer than 300um compared to neurons with processes longer than 200 um (**f**, data shown are average length of neurites independent t-test, p < 0.05).

Our ability to identify specific sub-populations of CSNs *in vitro* combined with a comprehensive understanding of the *in vivo* gene expression programs that are driving the post-natal patterning of the CST will allow for streamline targeting of robust pro-growth interventions.

## Discussion

In this study, we used a combination of viral retrograde labeling from the cervical and lumbar spinal cord, FACS and scRNAseq to annotate a comprehensive transcriptional atlas of the adult CST. Using a refined extraction, dissociation, and single cell sorting pipeline, we were able to maintain cytoplasmic integrity of large diameter pyramidal neurons for whole cell sequencing. The intersection of retro-labeling and scRNAseq revealed that *Wnt7b* is a selective marker for CSNs among other layer V neurons in M1, *Cacng7* is enriched FL CSNs, *Slc16a2* is enriched in HL CSNs, and *Rspo2* is enriched in a small subset of CSNs with collaterals in both the FL and HL. Additionally, we found that limb specific subsets of CSNs terminate differentially in the dorsoventral plane of spinal grey matter are also transcriptionally unique. Furthermore, we show that scRNAseq of layer V neurons at P5 showed that these novel CSN markers are expressed early postnatally. We confirmed these data with smFISH in dissociated E17 cortical neurons after 8 DIV. From these findings we can conclude that CSNs are molecularly unique among non-projection layer V neurons and can be further sub-divided based on the location of their rostrocaudal and dorsoventral terminal arbors. Leveraging this molecular heterogeneity provides a powerful opportunity to explore the transcriptional machinery that drives development, maintains homeostasis, and actuates reactive changes after trauma and disease within this critical motor pathway. Comprehensive transcriptional insight in these physiological and maladaptive processes will allow for refined therapeutic interventions to repair the damaged CST with spatial and temporal precision.

Previous work using scRNAseq of adult motor regions including M1 and anterior lateral motor cortex (ALM) established a taxonomy of neuronal and non-neuronal cell types and identified region-specific differences by comparing these cells to those in visual cortex.^15, 16, 26^ To describe the broad diversity observed between and among neuronal and non-neuronal cell types spanning the cortical depth in motor regions, cell-type specific markers describing pyramidal tract (PT), near-project (NP), corticothalamic (CT) and intratelencephalic (IT) neurons were identified.^26^ These data were confirmed by retrograde tracing from the pons and medulla and scRNAseq of ALM neurons further validating the PT designation.^16, 31^ However, these studies did not pursue tracing from the spinal cord, thus rendering a full PT atlas incomplete. Spinally traced PT neurons have been previously explored from a developmental perspective by combining retrograde tracing from the pons and upper cervical spinal cord early postnatally using temporal bulk microarray sequencing of FACS-purified neurons.^12^ Authors identified *Bcl11b, Crym,* and *Crim1* as CSN-specific markers up to P14, leaving the adult CSN transcriptome unexplored and spatially incomplete. Building on these studies, we sought to comprehensively profile spinally projecting neurons in the adult by performing retrograde tracing and scRNAseq from the adult CST. By leveraging the power of scRNAseq, we were able to identify and anatomically confirm robust CSN-specific and limb-specific genetic markers that can be differentially exploited for developmental, interventional, and functional analyses.

Functional subtypes among CSNs in M1 were previously described using a combination of slice electrophysiology and morphological analyses.^10^ Unbiased clustering based on these properties found that amongst CSNs, FL and HL-projecting neurons fired action potentials at the fastest rate and with the most regular pattern, and that burst firing was exclusive to cervical CSNs.^10^ This is consistent with FL-specific expression of *Cacng7,* a type II transmembrane AMPA receptor regulatory protein that enhances glutamate-evoked currents from GluR1.^39^ Molecular divergence between FL and HL CSNs revealed in our sequencing data supports these findings and underscores the need to approach these sub-branches of the CST independently when considering therapeutic interventions. This spinal enlargement level transcriptional resolution resolves broad FL and HL-level functional differences; however, there could also be within and between spinal level specificity. Intersegmental specificity was explored using *in vivo* calcium imaging from M1 during a pellet-retrieval task, showing that activity within traced CSNs originating from discrete spinal segments coordinates different phases of forelimb movement.^40^ These data would predict that transsynaptic retrograde tracing from FL muscles would further resolve functional sub-units within the cervical spinal cord differentially innervated by a subset of FL CSNs.

To explore within level specificity, we analyzed potential ligand receptor interactions in the lumbar spinal cord and found that HL CSNs synapsed upon SINs predominantly in the dorsal horn. Anterograde tracing studies supported this finding (**Fig. 4**). While a sensory modulation function for HL CSNs seems counterintuitive, a recent study showed that the lumbar CST functions exclusively to gate sensory input via primary afferent depolarization (PAD), and CST-mediated motor commands to the lumbar spinal cord are not direct, but rather use polysynaptic circuits in the upper cervical cord.^41^ Our data supports this finding as 3 of the 9 SIN populations that receive pre-synaptic input from HL CSNs are inhibitory, and thus central to PAD. Additionally, mice deficient in the HL CSN maker *Slc16a2* do not display gross motor abnormalities, but do exhibit shorter latency during a hot plate test, suggestive of hyperalgesia.^41, 42^ In further support, mice that underwent rotarod training showed extensive *cfos* labeling in both dorsal and ventral horns in the lumbar spinal cord confirming that motor output mediated in part by the CST engages both excitatory and inhibitory circuits.^13^

In addition to discrete FL and HL populations of CSNs, retrograde tracing shows that a fraction of CSNs have collaterals in both cervical and lumbar cord (**Fig. 1c)**. Transcriptional profiling of this dual projecting population shows that it is molecularly divergent from FL and HL CSNs and can be independently labeled via enriched expression of *Rspo2*. The existence and function of dual projecting CSNs is controversial, as extensive electrophysiological studies in the cat showed that stimulation of FL CSNs failed to elicit activity in the lumbar spinal cord and vice versa.^43^ However, the presence of dual labeled CSNs in our study and others, suggests that these neurons are either unique to the rodent or perhaps are functionally silent in the intact spinal cord, but may be pre-wired to assume an auxiliary role in the event of trauma or degeneration.^44–46^ Indeed, previous work from the Schwab lab has highlighted a potential functional role of these FL/HL CSNs after CNS trauma. They found an increase in functional sprouting of HL CSNs into the cervical spinal cord after stroke and SCI. ^47, 48^ These data could represent an activation of dual FL/HL CSNs, and furthermore suggesting that this sub-population has a high intrinsic growth capacity. Our data shows that this population of CSNs is enriched in *Rspo2*, leveraging transcriptional differences of *Rspo2*+ neurons after injury could identify novel pro-plasticity factors. Furthermore, specifically targeting *Rspo2*-expressing CSNs for intervention could enhance recovery after spinal cord injury.

As the CST enters the spinal cord and is wired entirely postnatally, the tract provides a unique opportunity to identify and ultimately exploit the molecular mechanisms that support different phases of CNS axon growth including initial outgrowth, scaling, pathfinding, co-lateral sprouting, target recognition, and functional synapse formation and refinement. Recapitulating developmental gene expression programs including targeting *Pten*, *Klf6*, *Sox11* after CNS injury stimulates robust axon regeneration, but limited functional recovery. These data suggest that additional factors delivered in a temporally specific manner are required to stimulate functional axon growth. Leveraging the CSN specific markers identified in this study can overcome these spatial and temporal challenges. Anterograde and retrograde tracing studies show that the CST reaches C8 by P2 but doesn’t enter the lumbar cord until P9.^2^ To date studies that have sought to explore CSN subtypes have used microarray sequencing of retrogradely traced CST axons from the cervical spinal cord at P6 and P14.^12^ Both time points would sample a mixed population of FL and HL CSNs and thus reveal an aggregate gene expression profile detailing a spectrum of growth associated mechanisms and phenotypic markers. Our data show that adult FL and HL CSN markers are preserved at P5. Transcriptional profiling at this time point provides insight into two unique developmental profiles: cervical-projecting CSNs that have reached their terminal location and are engaged in local plasticity, and lumbar-projecting CSNs that are still engaged in long-distance axon growth. Pathway analysis reveals that FL CSNs expressed genes associated with sprouting, consistent with these axons forming collaterals in grey matter at P5. While HL CSNs primarily expressed genes associated with axon guidance, consistent with these axons navigating structures along the rostrocaudal axis of the spinal cord at P5.

Data from our laboratory supports the notion that FL and HL CSNs require independent interventions to stimulate functional axon. In a recent study, we completed bulk transcriptional profiling of retrogradely traced CST axons that had undergone pyramidotomy (PyX)-induced functional sprouting. Differential gene expression analysis revealed that members of the LPAR1 interacting pathway and the 3-phosphoinositide degradation pathway were enriched in sprouting CSN. Targeting these pathways via viral over expression of *Lppr1* and *Inpp5k* in intact CSNs after contralateral PyX resulted in significant sprouting of intact CST axons into the denervated side of the spinal cord.^8, 9^ Assessment of fine motor skills showed that the lesioned FL performed better than the lesioned HL, suggesting that our treatments were having a greater impact in FL CSNs. These data support limb specific therapeutic interventions, as the pro-growth factors we identified in our original *in vivo* screen were identified via retrograde tracing from the cervical spinal cord. Future studies will reveal if tracing from the lumbar cord after contralateral PyX will reveal a unique set of pro-axon growth pathways.

Thus, exploiting the FL and HL specific CSN markers identified here will allow for unparalleled temporal precision to identify the mechanisms that drive all phases of CST axons growth and ultimately aid in the design of specific pro-growth therapeutics that target either whole subsets of the CST, i.e., all lumbar projecting axons after severe thoracic SCI, or more refined interventions that stimulating co-lateral sprouting after less severe partial injuries.

## Supplementary Figures

**Supplementary Figure 1.**
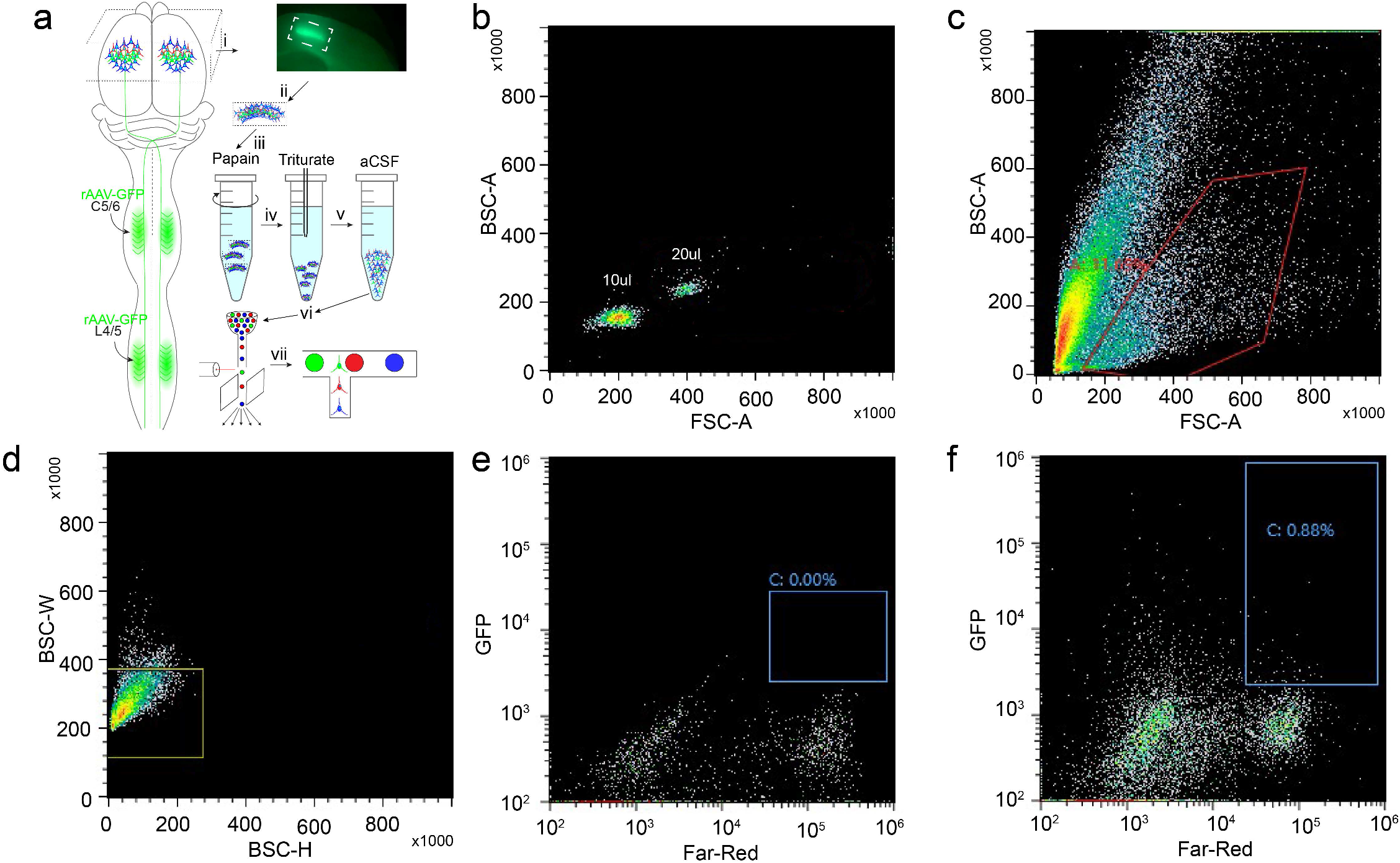
Schematic overview of the retroseq procedure from figure 1 (**a**). Mice received retrograde injections of rAAV-CAG-GFP into either C6/7 or L4/5 (i). After a 2-week incubation, M1 was macrodissected (ii), enzymatically dissociated in papain (iii), mechanically dissociated (iv), and incubated in aCSF (v). Traced, *Rbp4*-positive, and nonfluorescent cells were then sorted using FACS (vi) and sequenced using 10x (vii). FACS gating strategy (**b-f**). 10um and 20um beads were first sorted to set the lower limit for cell size (**b**). The first gate was then selected based on the beads, selecting for neurons with a soma size of at least 10um (**c**). Doublets were then removed by sorting based on side scatter width and height (**d**). To set the gate for GFP, a GFP-negative sample was sorted, and the gate was drawn so no non-fluorescent cells were selected (**e**). The traced samples were then sorted, and GFP-positive cells were collected for 10x sequencing (**f**).

**Supplementary Figure 2.**
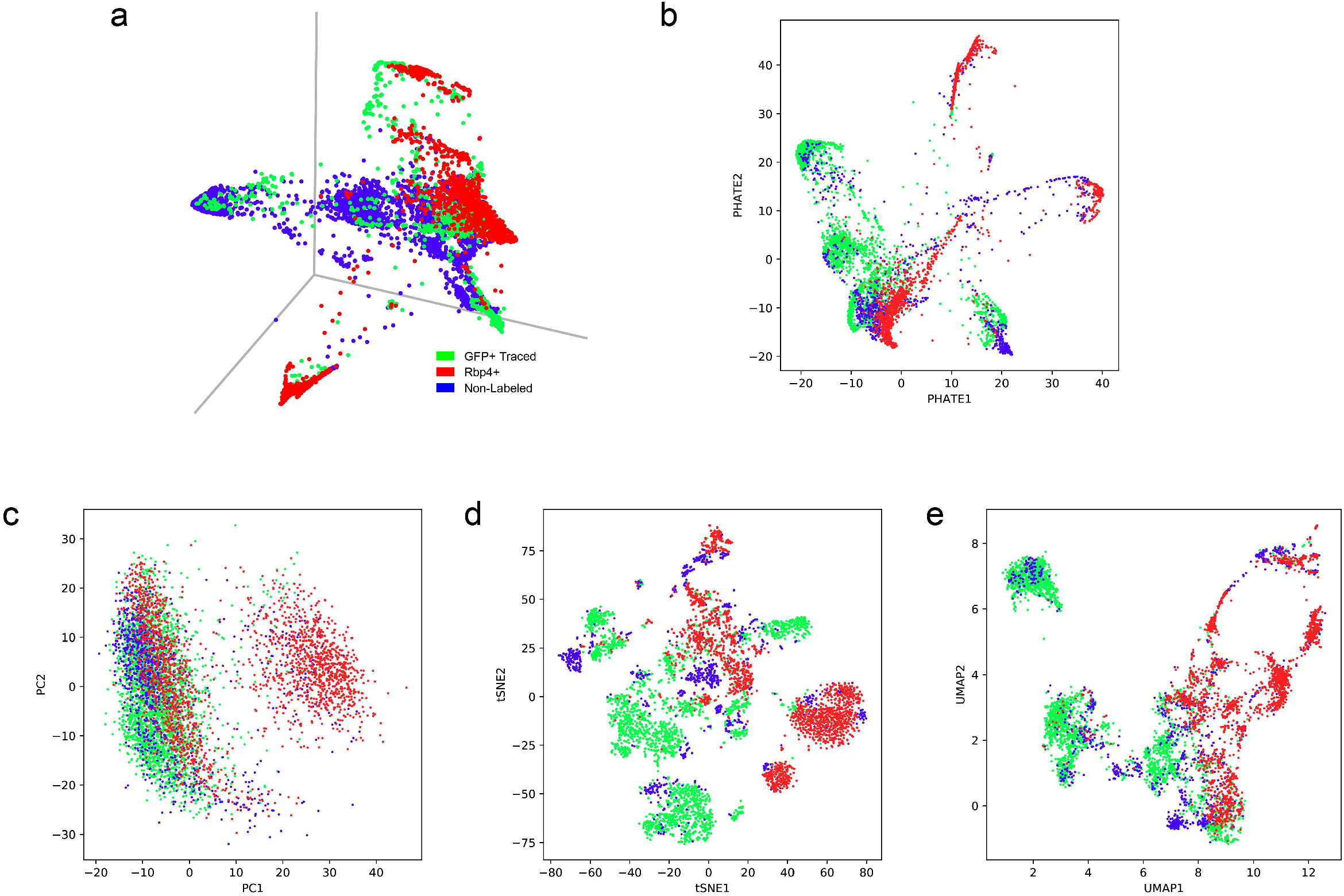
Comparison of dimensionality reduction visualization tools colored by cell identify (**a-e**). Three-dimensional PHATE (**a**), two-dimensional PHATE (**b**), principal component analysis (**c**), t-Distributed Stochastic Neighbor Embedding (t-SNE, **d**), and uMAP (**e**) show distribution of traced, *Rbp4*+ and non-fluorescent neurons.

**Supplementary Figure 3.**
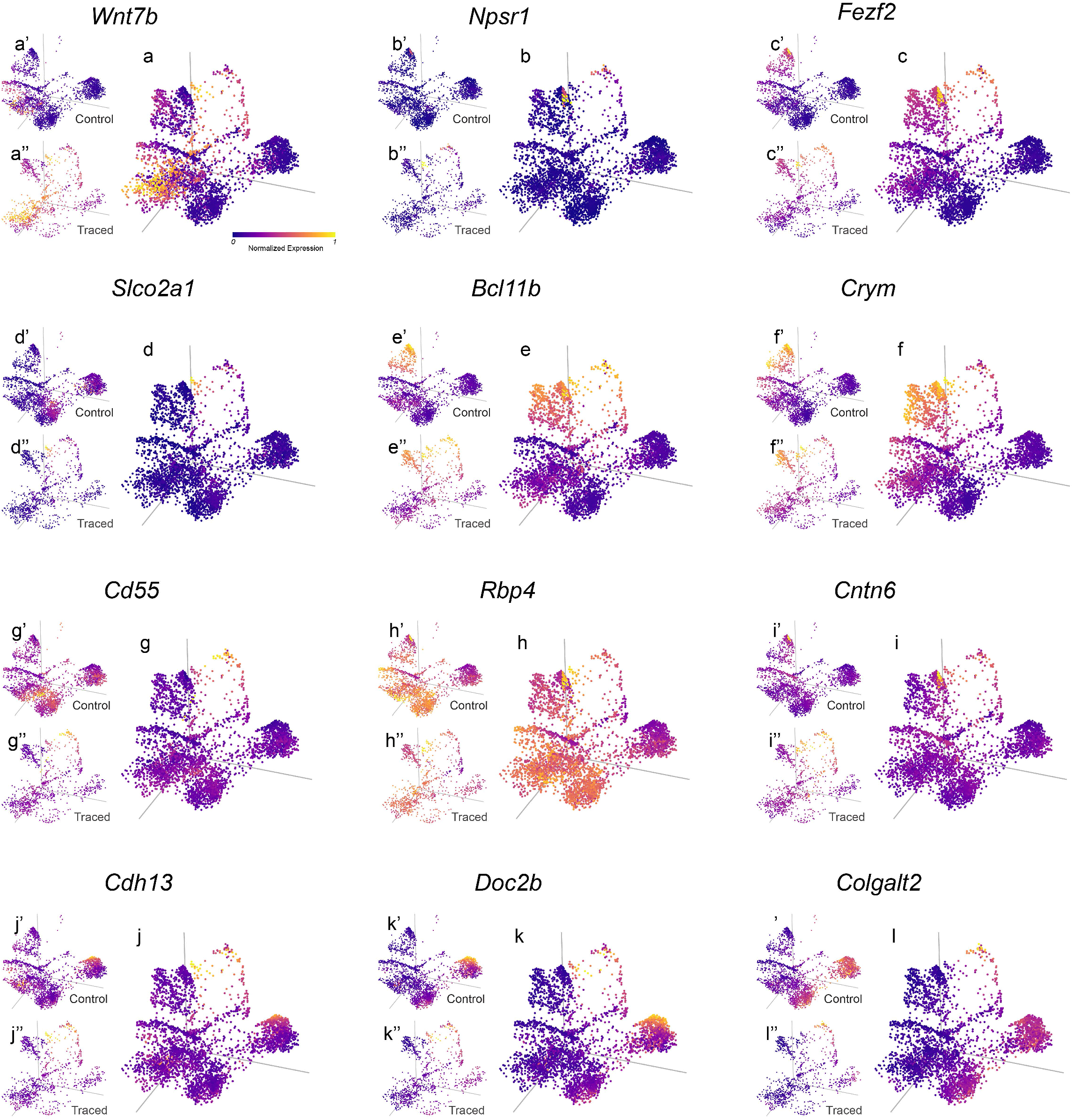
PHATE maps of traced and control CSNs, colored by normalized expression of *Wnt7b* in all cells (**a**), in control cells (**a’**) and in CSNs (**a”**), *Npsr1* in all cells (**b**), in control cells (**b’**) and in CSNs (**b”**), *Fezf2* in all cells (**c**), in control cells (**c’**) and in CSNs (**c”**), *Slco2a1* in all cells (**d**), in control cells (**d’**) and in CSNs (**d”**), *Bcl11b* in all cells (**e**), in control cells (**e’**) and in CSNs (**e”**), *Crym* in all cells (**f**), in control cells (**f’**) and in CSNs (**f”**), *Cd55* in all cells (**g**), in control cells (**g’**) and in CSNs (**g”**), *Rbp4* in all cells (**h**), in control cells (**h’**) and in CSNs (**h”**), *Cntn6* in all cells (**i**), in control cells (**i’**) and in CSNs (**i”**), *Cdh13* in all cells (**j**), in control cells (**j’**) and in CSNs (**j”**), *Doc2b* in all cells (**k**), in control cells (**k’**) and in CSNs (**k”**), and *Colgalt2* in all cells (**l**), in control cells (**l’**) and in CSNs (**l”**).

**Supplementary Figure 4.**
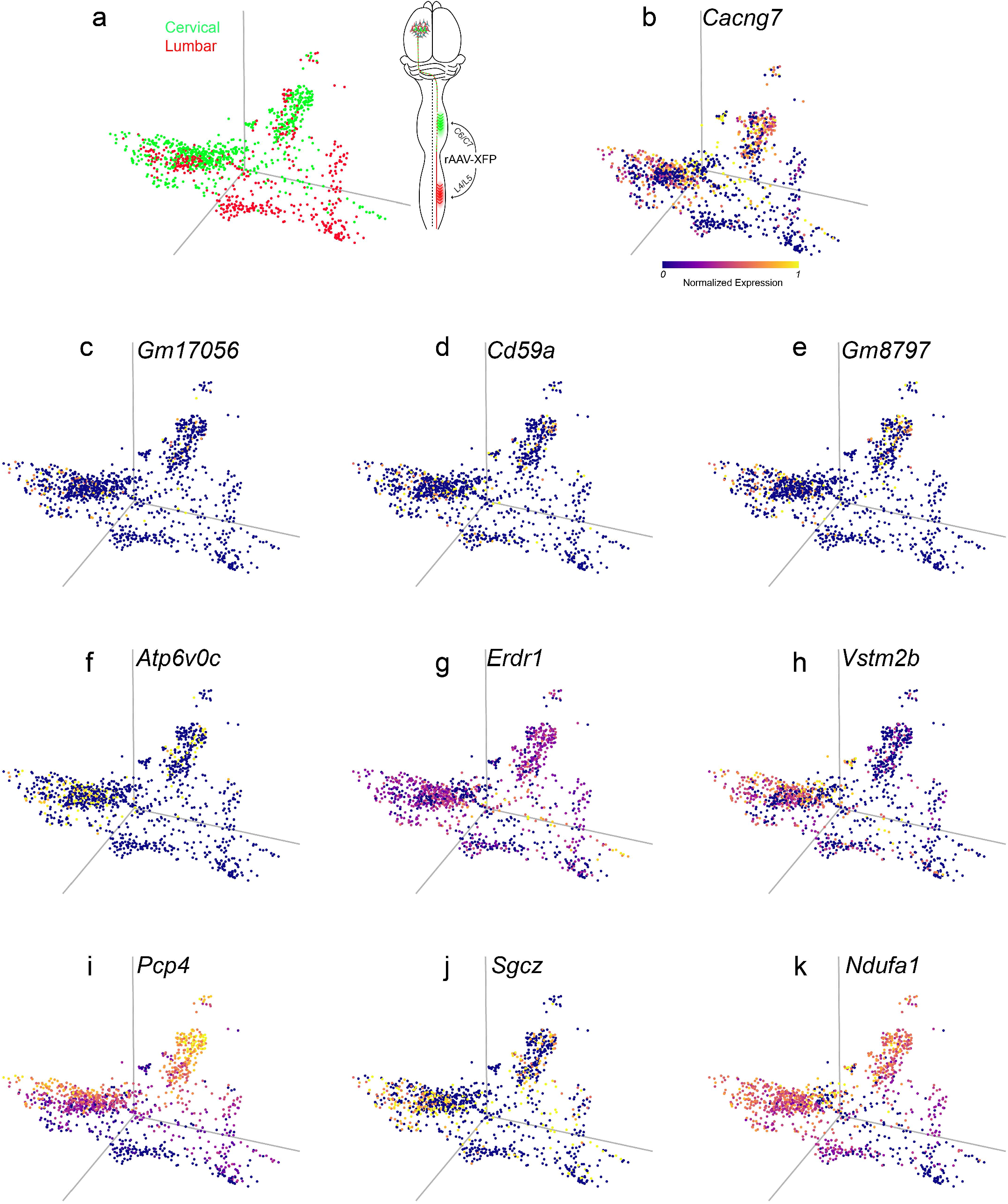
PHATE map of CSNs colored by FL and HL cell identity (**a**). PHATE maps colored by normalized expression of *Cacng7* (**b**), *Gm17056* (**c**), *Cd59a* (**d**), *Gm8797* (**e**), *Atp6v0c* (**f**), *Erdr1* (**g**), *Vstm2b* (**h**), *Pcp4* (**i**), *Sgcz* (**j**), *Ndufa1* (**k**).

**Supplementary Figure 5.**
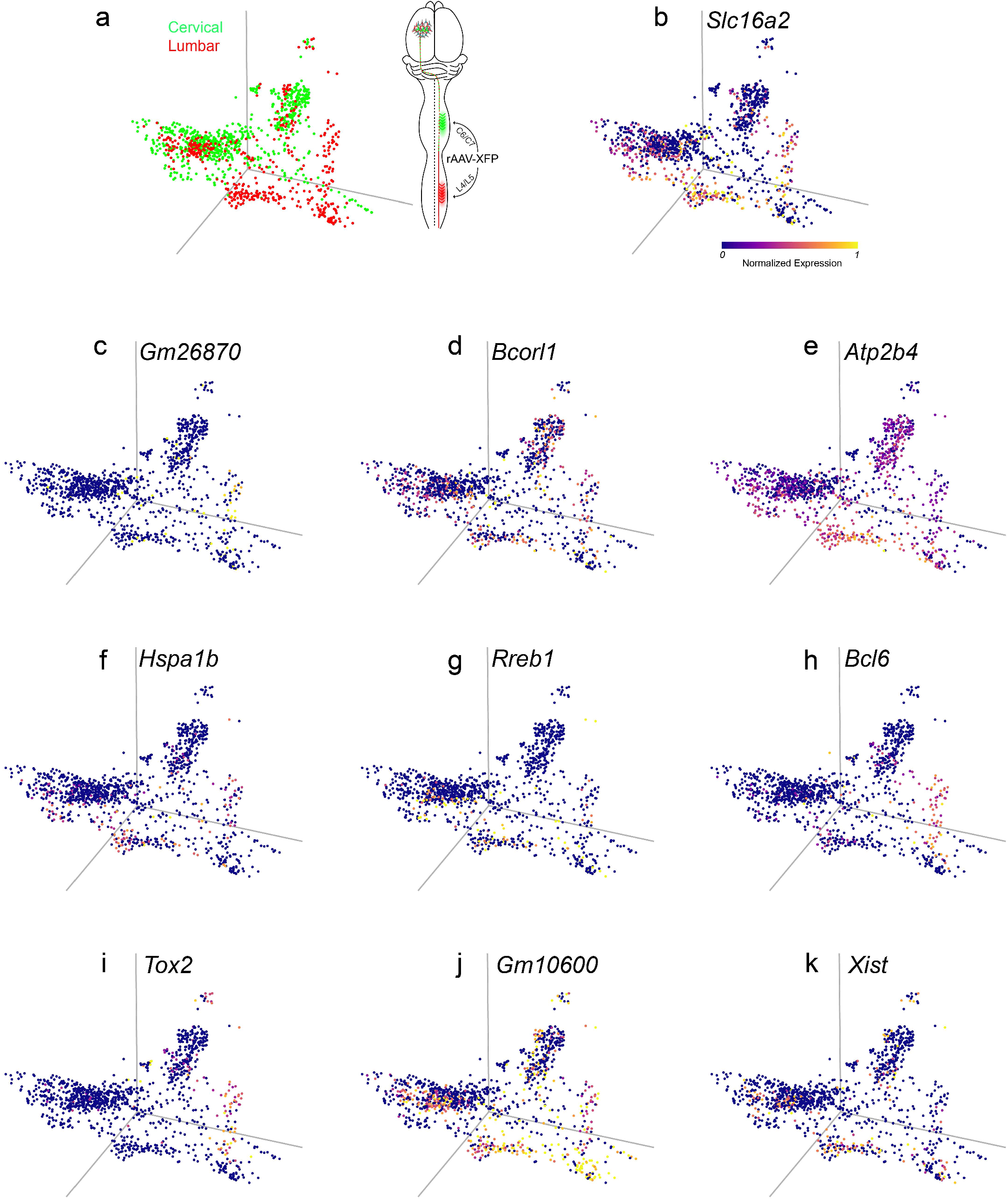
PHATE map of CSNs colored by FL and HL cell identity (**a**). PHATE maps colored by normalized expression of *Slc16a2* (**b**), *Gm26870* (**c**), *Bcorl1* (**d**), *Atp2b4* (**e**), *Hspa1b* (**f**), *Rreb1* (**g**), *Bcl6* (**h**), *Tox2* (**i**), *Gm10600* (**j**), *Xist* (**k**).

**Supplementary Figure 6.**
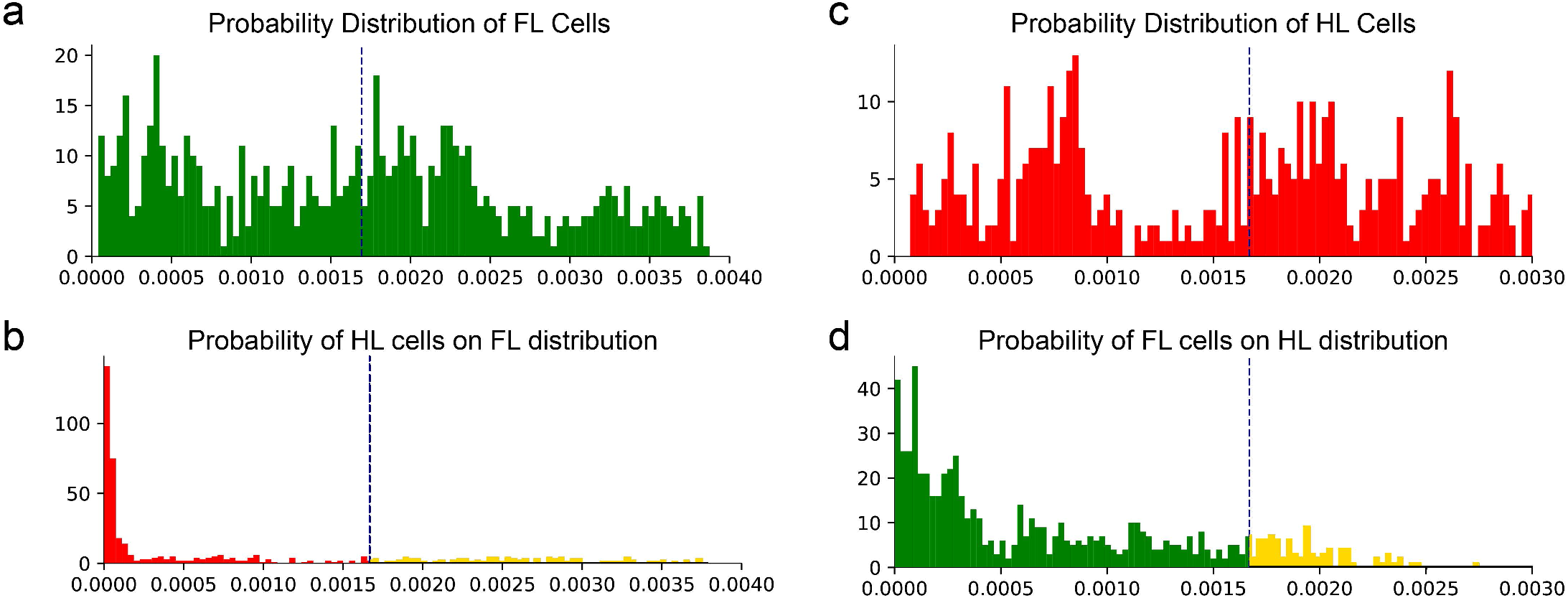
Likelihood histogram of FL cells based on kernel density estimate of FL CSNs (**a**). Likelihood histogram of HL cells based on kernel density estimate of FL CSNs (**b**). Likelihood histogram of HL cells based on kernel density estimate of HL CSNs (**c**). Likelihood histogram of FL cells based on kernel density estimate of HL CSNs (**d**).

## Methods

### Animals

*Rbp4* Cre mice (gift from Dr. David Berson, Brown University) were crossed with a cre-dependent tdTomato reporter line (Ai14, The Jackson Laboratory) to label deep layer cortical neurons. All sequencing experiments were conducted using Ai14 mice without the presence of cre. Fluorescent *in situ* hybridization experiments were conducted on C57BL/6J mice ordered from Jackson Laboratories.

### Surgery

All procedures and postoperative care were performed in accordance with the guidelines of the Institutional Animal Use and Care Committee (IACUC) at Yale University.

#### Intraspinal retrograde AAV injections for scRNAseq

To complete retrograde labeling of cervical-projecting CSNs, adult mice (n= 8, 4 males, 4 females) were anesthetized with ketamine (100mg/kg) and xylazine (15mg/kg) and placed in a stereotaxic frame (Stoelting, USA). An incision was made over the cervical enlargement and the C5-C8 vertebrae revealed by blunt dissection of overlying muscle. A bilateral laminectomy was performed to expose the underlying C5-C8 spinal cord, and a small incision was made in the dura mater. The tip of a pulled glass capillary tube attached to a Micro4 infusion device (World Precision Instruments, USA) was slowly inserted stereotaxically to a depth of 500 μm into the C5 level of the spinal cord and approximately 600 μm lateral from the midline. Thirty seconds after introduction of the capillary tube 100 nl of rAAV-CAG-GFP (Addgene, 37825-AAVrg) was infused into the spinal cord over 2 minutes. The tip was left in situ for an additional 30 seconds prior to removal. This procedure was completed 7 additional times bilaterally at the same coordinates at C6, C7 and C8 resulting in a total infusion of 800 nl. Muscle was sutured with Vicryl and skin with monofilament suture. All animals received post-surgical antibiotics (Ampicillin, 100mg/kg subcutaneously) and analgesia (Buprenorphine 0.05mg/kg subcutaneously) for 2 days post lesion. All animals recovered uneventfully.

To complete retrograde labeling of lumbar-projecting CSNs, adult mice (n=16, 8 males, 8 females) were anesthetized with ketamine (100mg/kg) and xylazine (15mg/kg) and placed in a custom-built spine stabilizer (Farrar, *et al*.) Using the last rib as a landmark, an incision was made and a bilateral laminectomy was performed to expose L4-L5 spinal cord. The tip of a pulled glass capillary tube attached to a Micro4 infusion device was slowly inserted stereotaxically to a depth of 500 μm into the L4 level of the spinal cord and approximately 500 μm lateral from the midline. Thirty seconds after introduction of the capillary tube 100 nl of rAAV-CAG-GFP was infused into the spinal cord over 2 minutes. The tip was left in situ for an additional 30 seconds prior to removal. This procedure was completed 7 additional times bilaterally at the same coordinates at L4 and L5 resulting in a total infusion of 800 nl. All animals received post-surgical antibiotics (Ampicillin, 100mg/kg subcutaneously) and analgesia (Buprenorphine 0.05mg/kg subcutaneously) for 2 days post lesion. All animals recovered uneventfully.

### Adult whole single-cell isolation

To isolate intact adult CSNs, mice that received retrograde injections into either the lumbar or cervical cord were anesthetized with isoflurane and transcardially perfused with artificial cerebrospinal fluid (aCSF) ten days after rAAV injection. The aCSF that consisted of CaCl2 (0.5 mM), glucose (25 mM), HCl (96 mM), HEPES (20 mM), MgSO4 (10 mM), NaH2PO4 (1.25 mM), myo-inositol (3 mM), N-acetylcysteine (12 mM), NMDG (96 mM), KCl (2.5 mM), NaHCO3 (25 mM), sodium L-ascorbate (5 mM), sodium pyruvate (3 mM), taurine (0.01 mM), thiourea (2 mM), trehalose (13.2mM) and was bubbled with carbogen gas (95% O2 and 5% CO2) ^1^. The brain was dissected and submerged in ice-cold bubbled aCSF for three minutes before being transferred to a brain matrix where 500um sections around M1 were sliced and sections were transferred to a petri dish containing ice cold bubbling aCSF. Regions of M1 containing labeled CSNs were dissected under a fluorescent microscope and transferred to a 5ml Eppendorf tube with dissociation buffer with papain.

The dissociation buffer contained sodium sulfate (82mM), potassium sulfate (30mM), HEPES (10mM), glucose (10mM) and magnesium chloride (5mM) ^2^. Thirty minutes prior to dissociation, the dissociation buffer was warmed to 34□ and 5 mL was added to a vial of lyophilized papain (Worthington Biochemical Corporation). The solution was diluted 1:2 with additional dissociation buffer prior to sample dissociation.

The sample in dissociation buffer with papain was incubated at 34□ for 70 minutes on a shaker at medium speed. After 70 minutes, the dissociation solution with papain was replaced with a Stop buffer containing 5 ml of dissociation buffer, 5mg Ovomucoid Protease Inhibitor (Worthington Biochemical Corporation) and 10mg bovine serum albumin for 5 minutes on ice. The Stop solution was then replaced with 800ul of dissociation buffer and triturated with Pasteur pipettes with decreasing diameters (600um, 300um and 150um) and placed on ice. To differentiate between intact neurons and nuclei, a cell-permeable dye was added at a concentration of 1:1000 for 5 minutes (invitrogen). The sample was then centrifuged at 300g for 10 minutes at 4□ and resuspended in 500ul of dissociation buffer.

### Neonatal whole single-cell isolation

To isolate intact neurons at P5, a modified protocol to the adult whole single-cell isolation was used. Briefly, *Rbp4* cre: Ai14 mice were transcardially perfused with aCSF and tdTomato-positive layer V neurons were used to guide macrodissection. Neurons were then enzymatically dissociated in papain solution for 20 minutes at 34□ on a shaker at medium speed. The neurons were then processed similarly to adult neurons.

### Fluorescence activated cell sorting (FACS)

Fluorescent single cells were isolated from *rbp4* cre: Ai14 mice and Ai14 mice injected with rAAV-CAG-GFP. Sorting was performed on the Sony SH800 with a 130um nozzle, a sheath pressure and sample pressure of less than 9 PSI. To ensure pyramidal neurons were being sorted, size reference beads were used to set a minimum diameter of 10um (Spherotech) for all collected cells. To exclude non-intact cells, the cells were sorted for the presence of the cell-permeable dye. A sample of non-fluorescent tissue from adjacent regions of the cortex were used as negative controls to set the gate for the presence of fluorescence (GFP or tdTomato). The sample was then sorted into Eppendorf tubes containing 4L dissociation buffer and immediately sequenced.

In total we used 28 animals (4 *rbp4* cre: Ai14, 8 Ai14 injected with rAAV-CAG-GFP into C6/7, 16 Ai14 injected with rAAV-CAG-GFP into L4/5). Dissection of layer V of M1 was guided by fluorescence and consistent across all animals.

### Characterization of cell morphology after dissociation and FACS

ImageStream data and images were acquired using the Amnis Inspire software and the Amnis ImageStreamX Mk II two-camera system, using the 405, 488, and 561 lasers and the 60× objective. Data were analyzed using Amnis IDEAS software (Luminex Corp.).

Additionally, cells were sorted into a single cell of a 96-well plate containing 4% paraformaldehyde (PFA) and imaged on an epifluorescent microscope (LeicaMicrosystems) for confirmation of intact morphology after FACS.

### 10X single-cell RNAseq

#### Construction of 10X Genomic Single Cell/Nuclei 3’ RNA-Seq libraries CITE-Seq Libraries and sequencing

The single cell protocol enables short read sequencing to deliver a scalable microfluidic platform for digital gene expression of 500-10,000 individual cells per sample.

#### Sample Preparation

The first step of scRNA-Seq involves preparation of the single cell suspension, which would differ depending on the starting sample types. With samples that would perform CITE-Seq (Cellular Indexing of Transcriptomes and Epitopes by Sequencing) together with transcriptomics analysis, the cells are then bound to the oligonucleotide barcoded antibodies of surface proteins. Cell surface proteins can be labeled using a specific protein binding molecule, such as an antibody conjugated to a Feature Barcode oligonucleotide. Additional nuclei isolation would be needed for single nuclei transcriptomics analysis.

#### GEM Generation and Barcoding

Single cell suspension in RT Master Mix is loaded on the Single Cell A Chip and partition with a pool of about 750,000 barcoded gel beads to form nanoliter-scale Gel Beads-In-Emulsions (GEMs). With each gel bead, it has primers containing (i) an Illumina® R1 sequence (read 1 sequencing primer), (ii) a 16 nt 10x Barcode, (iii) a 12 nt Unique Molecular Identifier (UMI), and (iv) a poly-dT primer sequence (30nt). Upon dissolution of the Gel Beads in a GEM, the primers are released and mixed with cell lysate and Master Mix. Incubation of the GEMs then produces barcoded, full-length cDNA from poly-adenylated mRNA.

#### Post GEM-RT Cleanup, cDNA Amplification and library construction

Silane magnetic beads are used to remove leftover biochemical reagents and primers from the post GEM reaction mixture. Full-length, barcoded cDNA is then amplified by PCR to generate sufficient mass for library construction. Enzymatic Fragmentation and Size Selection are used to optimize the cDNA amplicon size prior to library construction. R1 (read 1 primer sequence) are added to the molecules during GEM incubation. P5, P7, a sample index, and R2 (read 2 primer sequence) are added during library construction via End Repair, A-tailing, Adaptor Ligation, and PCR. The final libraries contain the P5 and P7 primers used in Illumina bridge amplification.

#### Sequencing libraries

The Single Cell 3’ Protocol produces Illumina-ready sequencing libraries. A Single Cell 3’ Library comprises standard Illumina paired-end constructs which begin and end with P5 and P7. The Single Cell 3’ 16 bp 10x Barcode and 12 bp UMI are encoded in Read 1, while Read 2 is used to sequence the cDNA fragment (91bp). Minimum sequencing depth is 20,000 read pairs per cell.

### Single-cell RNAseq analysis

#### Quality Control and Normalization

Single-cell analyses were performed with Python 3.8.5 and the scipy (1.5.2) and numpy (1.19.2) modules. After extracting the single-cell gene expression matrix, we used the scprep (1.0.3) package in python to eliminate non-neurons and normalize the data^3^. Briefly, we filtered out cells that had a library size of less than 2500 UMIs as well as cells that had mitochondrial gene expression in the top tenth percentile. After a library size normalization, we used the python package MAGIC to impute missing data ^4^. Using the MAGIC output gene expression matrix, we filtered out contaminating cell types (endothelial cells, microglia, astrocytes) as well as inhibitory interneurons, resulting in an approximately 1% reduction from the starting neuronal population. We then square-root transformed the data for further downstream analysis.

#### Distinguishing CSNs from adjacent L5 neurons

To identify CSN-specific markers, we first needed to identify putative GFP-negative cells from the samples of *Rbp4* and non-fluorescent sequenced neurons. Due to inherent inefficiencies in viral labeling and FACS we utilized a computational approach. We approximated the distribution for each of our cell populations (*rbp4*, nonfluorescent, CSN, thalamic-projecting CSNs)^5^ using a gaussian kernel density estimate (KDE). Previous work has shown that *rbp4*+ cells do not have collaterals in the thalamus ^6^. Based on this finding we calculated the likelihood of each rbp4+ neuron with respect to the estimated distribution of thalamic cells. We then removed from analysis putative non-spinal rbp4+ cells, defined conservatively as those rbp4+ cells highly likely to have been generated from the thalamic distribution (likelihood greater than 95% of thalamic traced cells on the thalamic distribution). To identify putative non-spinal non-fluorescent neurons, we calculated the likelihood of each non-fluorescent cell for the GFP+ distribution. As above we removed non-spinal non-fluorescent cells from analysis as putative GFP+ cells defined as those found to have a GFP+ likelihood greater than 95% of GFP+ cells.

#### Identification of dual-cervical and lumbar projecting CSNs

To identify dual-projecting CSNs, we used KDE to approximate the FL and HL distributions. We then estimated the likelihood for each of our FL cells being generated from the distribution of traced HL cells and the same for HL cells on the distribution of traced FL cells. We identified putative dual-projecting CSNs as those traced from one region that had a likelihood measure from the other distribution greater than the median traced cell from that distribution.

#### Differential expression analysis

Once we identified a pure non-spinally projecting CSN cell population (non-spinal *rbp4*+ and non-spinal nonfluorescent), we used the python package diffxpy to run a wilcoxon rank-sum test with a Benjamini-Hochberg correction for multiple comparisons to identify differentially expressed genes between GFP+ CSNs and non-spinal layer V cells^7^. Using the significantly differentially expressed genes (q < 0.05), we used recursive feature elimination to identify the top ten genes that best differentiate between CSNs and non-CSNs ^8^. To further validate these genes, we trained a Support Vector Machine classifier, with a RBF kernel, (SVM) to predict cell origin based on the top genes reserving half of our cells for cross validation ^8^. We trained this classifier over 100 random splits of the data and evaluated the accuracy, precision and recall of the SVM on the reserved test data. Differentially expressed genes were then used downstream in Ingenuity Pathway Analysis for identification of commonly regulated pathways. To identify FL and HL specific markers, a similar approach was taken. Briefly we used differential expression analysis followed by a linear SVM to confirm specificity.

#### CSN visualization

For low-dimensional visualization of all scRNAseq data, we used the python package PHATE in three dimensions, with the default settings ^4^.

#### Analysis of P5 CSNs

To select P5 CSNs, only *Wnt7b-*positive neurons were included in the analysis. Using the ‘cluster’ feature of the PHATE package, we performed k-means clustering on the embedded cellular positions, identifying two unique clusters corresponding with FL and HL CSNs. We then ran a wilcoxon rank-sum test with a Benjamini-Hochberg correction on the FL P5 CSNs against the FL P56 CSNs and on the HL P5 CSNs against the HL P56 CSNs. Differentially upregulated genes from both analyses were included in the Ingenuity pathway analysis.

#### Ligand-Receptor Analysis

To identify post-synaptic partners for lumbar corticospinal neurons (CSNs), cell:cell interactions were predicted using similar methods from Ximerakis et al.^9^ Briefly, the CellCellInteractions (CCI) dataset was used, which contained curated ligand/receptor pairs. This dataset was narrowed down to ligands and receptors that were significantly differentially upregulated in traced CSNs (compared to non-spinally projecting layer V neurons) and whose binding partners were expressed in lumbar interneurons from Sathyamurthy et al, ^10^. An expression index was calculated per lumbar interneuron cluster by dividing the number of genes from the narrowed down CCI dataset that were expressed by the total number of genes expressed and a one-tailed permutation test was used to calculate the significance of the expression index for each cluster. To further classify interactions unique to this circuit, a differential gene expression analysis using a Wilcoxon rank sum test between the nine identified lumbar interneuron clusters and the remaining clusters was used. Ligand/receptor pairs that were differentially upregulated in both the HL-CSNs and the lumbar interneurons were identified.

#### Injections for validation of the ligand-receptor analysis

For confirmation studies, mice received unilateral infusion of AAV-CAG-mCherry into motor cortex (M1). AAV transduction of CSNs was completed as previously described ^11^. Briefly, burr holes were made over sensorimotor cortex using a dental drill and 5 micro infusions of 150 nl of AAV-CAG-mCherry were made to a depth of 0.7 mm (co-ordinates, +1 mm to −1 mm posterior to bregma and 0.5 - 1.5 mm lateral to bregma) using a Hamilton syringe and Micro4 infusion device to deliver a total volume of 375 nl of each virus (750 nl total).

### Immunohistochemistry and smFISH Validation

#### Injections for in situ hybridization validation of scRNAseq

For confirmation studies, mice received bilateral injections of rAAV-CAG-GFP into the cervical cord and rAAV-CAG-tdTomato into the lumbar cord as described above for a total of 16 spinal injections. All animals received post-surgical antibiotics (Ampicillin, 100mg/kg subcutaneously) and analgesia (Buprenorphine 0.05mg/kg subcutaneously) for 2 days post lesion. All animals recovered uneventfully.

#### smFISH of adult cortical tissue

Ten days after retrograde AAV injections, mice were euthanized with isoflurane and transcardially perfused with 0.9% NaCl followed by 4% paraformaldehyde in PBS. Brains and spinal cords were dissected and post-fixed in 4% PFA overnight at 4°C. The next day, 15um thick sections were cut using a vibratome, mounted on superfrost plus slides and processed for smFISH according to the ACD RNAScope fluorescent protocol. Briefly, sections were post-fixed in 4% PFA and stained with a single probe. Each experiment included a positive and negative control probe to ensure validity. Prior to coverslipping, Immunofluorescence utilizing primary antibodies directed against green fluorescent protein (GFP, 1:2000, Abcam, USA), and mCherry (1:2000, Abcam, USA) and detected with secondary antibodies Alexa Fluor-488 and - 568, (1:500, 1:500 Life Technologies, Grand Island, NY) was used to visualized traced CSNs.

#### smFISH of adult cortical tissue- analysis

All analyses were performed by an experimenter blinded to probe identity. For quantification of smFISH results, images were taken at 40X on Leica Sp8 confocal microscope (LeicaMicrosystems). 3 images were taken per hemisphere, 3 sections were taken per animal and 3 animals were used for the analysis of each probe. For analysis, each image was separated into individual channels. The images of the traced neurons were then exported to photoshop, where the quick selection tool was used to select cell body regions of interest (ROI) and create masks. This was done manually as CSN axons can be very thick and automatic ROI selection often cannot distinguish between CSN axons and cell bodies. Images of the probe and the ROI masks are then uploaded to CellProfiler and puncta per cell are automatically calculated according to the CellProfiler manual ^12^.

#### Embryonic Cortical Neuron Cultures

Embryonic cortical neurons were cultured as previously described ^13^. Briefly, E17 mouse embryos were dissected from timed-pregnant C57Bl/6 mice and cortices were rapidly dissected in ice-cold Hibernate E medium (Brain Bits). Cortices were then digested for 30 minutes at 37°C using a digestion medium containing papain (25 U/ml, Worthington Biochemical), DNAse I (2000 U/ml, Roche), 2.5 mM EDTA, and 1.5 mM CaCl2 in a neuronal culture medium composed of Neurobasal A (Life Technologies) with B27 supplement (Life Technologies), 1% sodium pyruvate (Life Technologies), 1% glutamax (Life Technologies), and 1% penstrep (Life Technologies). After digestion, cortices were washed twice with the neuronal culture medium and triturated in 2 ml of medium. Triturated cells were passed through a 40 μm cell strainer (Corning) and counted. Neurons were then transferred to pre-warmed neuronal media and plated at a density of 2,000 cells per well in an 8-well glass Lab-Tek slide (Thermo Scientific) pre-coated with poly-D-lysine (Corning) and laminin (Life Sciences). Cells were cultured 8 days at 37°C prior to fixation.

#### smFISH of embryonic cortical cultures

After 8 DIV, neurons were fixed with 4% EM-PFA 30% sucrose diluted in PHEM buffer (300mM PIPES, 125 mM HEPES, 50 mM EGTAm 10 mM MgCl2) for 20 minutes and processed for smFISH according to the ACD RNAScope Cultured Adherent Cell Sample Preparation protocol. Briefly, after fixation, neurons were immediately dehydrated and stored in 100% ethanol at −20 °C until all samples were ready for staining to avoid batch effects. Slides were then rehydrated and stained with a single probe. Each experiment included a positive and negative control probe to ensure validity. Prior to coverslipping, Immunofluorescence utilizing primary antibodies directed against βIII-tubulin (1:1000, Promega) and secondary antibodies Alexa Fluor-488 (1:500, Abcam) were used to visualize cell processes.

#### smFISH of embryonic cortical cultures- analysis

All analyses were performed by an experimenter blinded to probe identity. Slides were imaged at 20X using an epifluorescent microscope (LeicaMicrosystems). Five images from 3 different wells from each biological n (2) were taken of individual neurons stained for each probe. The length of the longest neurite (from soma to tip of axon) was measured using the Simple Neurite Tracer (SNT) ImageJ plugin. smFISH puncta were counted manually.

